# FRET-GP – A Local Measure of the Impact of Transmembrane Peptide on Lipids

**DOI:** 10.1101/2023.08.26.554931

**Authors:** Garima C. N. Thakur, Arunima Uday, Piotr Jurkiewicz

**Affiliations:** J. Heyrovský Institute of Physical Chemistry of the Academy of Sciences of the Czech Republic, v.v.i., Prague, Czech Republic

## Abstract

Reconstitution of a transmembrane protein in model lipid systems allows studying its structure and dynamics in isolation from the complexity of the natural environment. This approach also provides a well-defined environment for studying the interactions of the protein with lipids. In this work we describe the FRET-GP method, which utilizes Förster resonance energy transfer (FRET) to specifically probe nanoenvironment of a transmembrane domain. The tryptophan residues flanking this domain act as efficient FRET donors, while Laurdan acts as acceptor. The fluorescence of this solvatochromic probe, is quantified using generalized polarization (GP) to reports on lipid fluidity in the vicinity of the transmembrane domain. We applied FRET-GP to study the transmembrane peptide WALP incorporated in liposomes. We found that the direct excitation of Laurdan to its second singlet state strongly contribute to GP values measured in FRET conditions. Removal of this parasitic contribution was essential for proper determination of *GP*_FRET_ – the local analogue of classical *GP* parameter. The presence of WALP significantly increased both parameters, but the local effects were considerably stronger (*GP*_FRET_ ≫ *GP*). We conclude that WALP restricts lipid movement in its vicinity, inducing lateral inhomogeneity in membrane fluidity. WALP was also found to influence lipid phase transition. Our findings demonstrated that FRET-GP simultaneously provides local and global results, thereby increasing comprehensibility of the measurement. We highlight the simplicity and sensitivity of the method, but also discuss its potential and limitations in studying protein-lipid interactions.

**TOC graphics:** 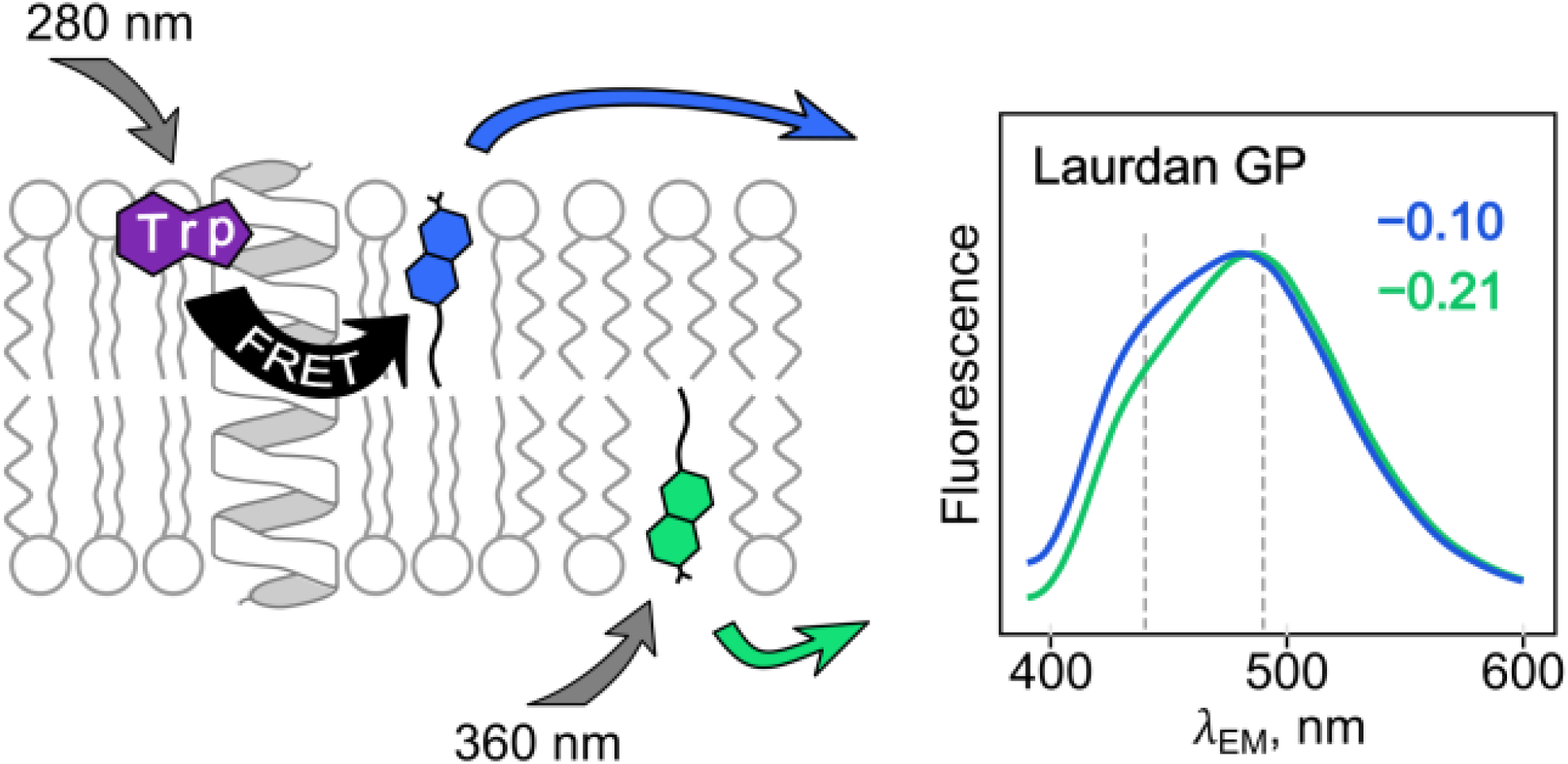

## INTRODUCTION

To understand structural and functional properties of a membrane protein, it is of great advantage if we can study them in isolation from myriads of other proteins present in a living cell. At the same time, maintaining a proper environment is crucial for preserving protein properties in vitro. This is especially true for membrane proteins, the crystallization of which outside lipid membranes usually leads to misfolding and aggregation.^1^ However, preserving or reconstructing their lipid environment is by no means trivial. This is likely the reason why only ∼1500 structures of integral membrane proteins have been identified so far,^2^ which is less than 1% of all known protein structures.^3^ This is despite the fact that membrane proteins make up ∼1/4 of all the proteins encoded in our genes^4^ and account for ∼60% of current drug targets.^5^

Lipid bilayer a complex and dynamic assembly of many lipid species^6^ and not a passive featureless 2D fluid that solvates integral membrane proteins. Instead, many protein and lipid are known to interact in a more or less specific way.^7, 8^ Importantly, these interactions can modulate the functions of the interacting proteins and also other proteins that are sensitive to the mechanical properties of the lipid membrane, e.g., mechanosensitive ion channels^9^. This is why reconstitution of proteins in lipid systems seems more appropriate than solubilizing them using detergents,^10^ allowing studying their properties in more details including their folding,^11^ or designing whole assemblies in the form of the so-called artificial cells^12^. As recently pointed out, proper choice of the lipid system might be sometimes even more important for transmembrane peptides, the size of which is comparable to that of a lipid.^13^

Liposomes with reconstituted proteins – proteoliposomes – are known from decades, but over the past 30 years we have seen continuous development of different models including planar systems,^14^ bicelles,^15, 16^ and nanodiscs^17^. Modern protocols utilize fusion of oppositely charged proteoliposomes,^18^ fusion of proteoliposomes with cushioned planar model membranes^19^ or even the stabilization of proteoliposomes with an exoskeleton of zeolitic-imidazole framework^20^. These systems can be successfully applied to study protein structure using modern NMR techniques,^21, 22^ but also cryogenic electron microscopy (cryo-EM) capable of single particle analysis^23, 24^. Experimental apparatus for studying protein-lipid interactions that are often weak and transient is constantly being developed, but we still face many challenges.^25^ Among them is certainly reproducibility of the formulations of the model systems. The strongly hydrophobic character of the typical transmembrane domain of a protein requires special handling during sample preparation to prevent their segregation from lipids. The same is true for transmembrane peptides. The resulting protein/peptide-to-lipid ratio can be broadly distributed even within a single sample.^26^

Here we present a simple fluorescence spectroscopy method that can be used to study the interactions of proteins or peptides with lipids in proteoliposomes and similar systems. It is based on the energy transfer from tryptophan to a solvatochromic probe Laurdan.^27^ Application of Förster resonance energy transfer (FRET) from the intrinsic fluorophore of a protein to the lipid probe provides a reliable test of the proximity of the two. Moreover, Laurdan fluorescence analyzed using generalize polarization method (GP)^28^ characterizes lipid mobility, which we showed to be affected by the presence of transmembrane domains.^29^ After overcoming the major obstacle of concurrent excitation of Laurdan to its second singlet state, the FRET-GP method provides unique insight into the lipids in the immediate vicinity of the peptide. Given the method is swift and only requires a standard spectrofluorometer, we believe it is also well-suited for routine monitoring of proteoliposomes.

## MATERIALS AND METHODS

### Materials

Cholesterol, 1-palmitoyl-2-oleoyl-*sn*-glycero-3-phosphocholine (POPC), and 1,2-dimyristoyl-*sn*-glycero-3-phosphocholine (DMPC) were obtained from Avanti Polar Lipids, Inc. (Alabaster, AL). 6-lauroyl-2-dimethylaminonaphthalene (Laurdan) was obtained from Molecular Probes (Eugene, OR, USA). HEPES was obtained from Fluka (Buchs, Switzerland). Ethylenediaminetetraacetic acid (EDTA) and calcium chloride (CaCl_2_) with purity ≥99% were purchased from Sigma-Aldrich (St. Louis, MO), and potassium chloride (KCl) with purity ≥98% from Fluka (Buchs, Switzerland). Salts and ethylenediamine tetraacetic acid (EDTA) were dissolved in Milli-Q (Millipore, USA) water. pH of EDTA solution was adjusted to 7.0 using KOH. 2, 2, 2-trifluoroethanol (TFE) was obtained from Alfa Aesar (Karlsruhe, Germany). WALP peptide (GTSTSKKWWLALALALALALALALALWWKKFSTS) with purity 89.1% was custom synthesized by Bachem Ltd (UK). Solutions of peptides were freshly prepared in TFE for each experiment. Organic solvents of spectroscopic grade were supplied by Merck (Darmstadt, Germany). All chemicals were used without further purification.

### Instrumentation

All measurements were performed in quartz spectroscopic cuvettes at 25°C or 37°C. A time-correlated single-photon counting (TCSPC) spectrometer 5000 U SPC, equipped with a 370 nm NanoLED 11 laser diode, >399 nm emission longpass filter, and a cooled Hamamatsu R3809U-50 microchannel plate photomultiplier (IBH, Glasgow, UK) was used to measure fluorescence decays of Laurdan. The width of the instrument response function measured for Ludox dispersion was 80 ps. Steady-state fluorescence was measured on FS5 spectrofluorometer (Edinburgh Instruments, Livingstone, UK). A Shimadzu UV-1601 spectrophotometer was used to measure UV-Vis extinction and a Zetasizer Nano ZS (Malvern Instrument, Worcestershire, UK) to measure peptidoliposome and liposome size distributions.

### Fluorescence Measurements

Five steady-state spectra were measured for each sample using the following set of excitation and emission wavelengths (*λ*_EX_, *λ*_EM_, respectively). Emission spectra of Laurdan and tryptophan were measured at *λ*_EX_=373 nm and *λ*_EX_=280 nm, obtaining emission spectra at 390–600 nm and 290–600 nm, respectively. Two excitation spectra (*λ*_EX_=200–430 nm) were measured for *λ*_EM_=440 nm and *λ*_EM_=490 nm to calculate GP and FRET-GP spectra. The excitation spectrum of tryptophan was measured for *λ*_EX_=200–320 nm with *λ*_EM_=330 nm. All spectra were measured with 1 nm step, 0.2 s per point in triplicate. Fluorescence decays of Laurdan were collected at *λ*_EX_=373 nm (pulsed laser diode) and *λ*_EM_=400–540 with 10 nm steps. A long pass filter >399 nm was used to minimize Rayleigh scattering of vesicle dispersions. A time-dependent fluorescence shift method (TDFS) was used to analyze the time-resolved data as previously described.^29^ More information about TDFS data analysis is given in the Supporting Information.

### Peptidoliposome Formulation

Appropriate volumes of chloroform solutions of POPC (or DMPC) and cholesterol, TFE solution of peptide, and methanol solution of Laurdan were mixed in a glass tube and dried under a nitrogen stream while continuously incubated in water bath (at ambient temperature) and then in vacuum for at least 4 h. Dry lipid/peptide film was hydrated in 1.0 mL of 10 mM HEPES buffer (pH 7.4, 140 mM KCl, 0.1 mM EDTA) and vortexed for 10 min.

Samples were then sonicated using a tip ultrasonicator (Sonopuls HD 2070, Bandelin electronic, Germany). Samples without peptide were prepared in the same way. The final lipid concentration was always 0.2 mM.

## RESULTS AND DISCUSSION

### Basic Characteristics of Peptidoliposomes

In this work we present a FRET-GP method that can be used to investigate to properties of proteoliposomes and similar systems, as well as to study protein-lipid interactions in lipid membranes in general. To demonstrate its potential, we characterize liposomes containing the transmembrane WALP peptide. We refer to these liposomes as peptidoliposomes to distinguish them from proteoliposomes, which incorporate whole membrane proteins. Our peptide belongs to WALP family^30^ and features alternating alanine-leucine hydrophobic core linked by tryptophan residues to hydrophilic flanking residues: GTSTSKKWWLALALALALALALALALWWKKFSTS. The presence of tryptophan is essential for the FRET-GP method, and it is also valuable for the precise determination of the peptide content.

Peptidoliposomes were formed by mixing all components in organic solvents, thoroughly drying the mixture, hydrating it with a buffer and sonicating. Chemical structures of the components and the scheme of the preparation procedure are shown in Figure 1A.

**Figure 1:**
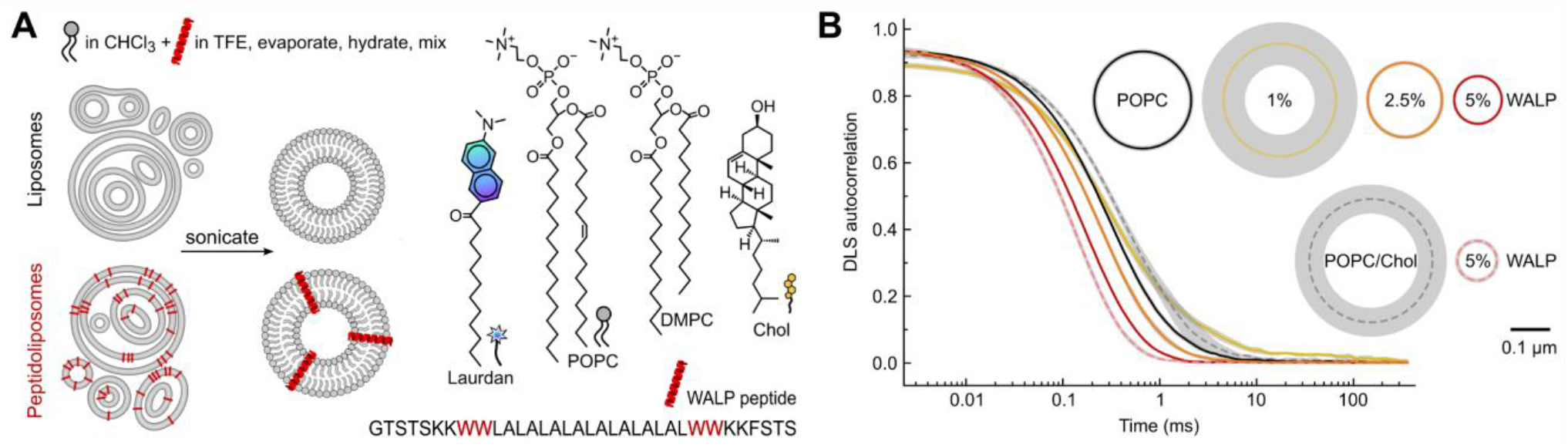
Vesicles formulation and sizes. (A) Scheme of peptidoliposome and liposome formulation, amino acid sequence of WALP peptide and chemical structures of POPC and DMPC lipids, cholesterol, and Laurdan fluorescent probe. (B) Vesicles sizes measured using dynamic light scattering. Autocorrelation curves are shown together with the mean hydrodynamic diameter of vesicles calculated using cumulant analysis. Vesicles are grouped based on lipid composition: POPC – upper row, POPC/Chol – lower row, and increasing WALP content – left to right; drawn to scale (scale bar). Gray areas represent standard deviation; please note that for some samples they are thinner than the line width.

We selected sonication over extrusion as it better preserves the molar ratios of all the sample components. Extrusion often causes peptide loss due to its interaction with extruder membrane. As determined by dynamic light scattering (DLS) and illustrated in Figure 1B, average diameters of our vesicles were between 0.05–0.30 μm, and no signs of aggregation nor precipitation were noticed. Peptide presence decreased vesicle sizes suggesting alteration of mechanical properties of the membrane. DLS results are analyzed in detail in the Supporting Information.

The absorbance of tryptophan is a more accurate measure of protein concentration than the absorbance of the peptide bonds alone^31^. Nevertheless, the turbidity of liposomal samples and the short light wavelengths significantly increase the contribution of Rayleigh scattering to the measured extinction spectra (Figure 2A). We removed this scattering by subtracting extinction of blank samples, which were vesicles without a peptide. Since the scattering in different samples differed, we used a simple fitting to achieve this step. The fitted blank spectra for 2.5% and 5% WALP samples are depicted as dotted lines in Figure 2A, and the resulting absorbance spectra are shown in Figure 2B. They are slightly red-shifted when compared to the absorbance of WALP dissolved in TFE and tryptophan dissolved in water. The content of Laurdan was determined in a similar manner, except that its presence does not affect the scattering, thus no fitting was necessary. The lower panel of Figure 2A shows the resulting differential Laurdan absorbance spectra.

**Figure 2:**
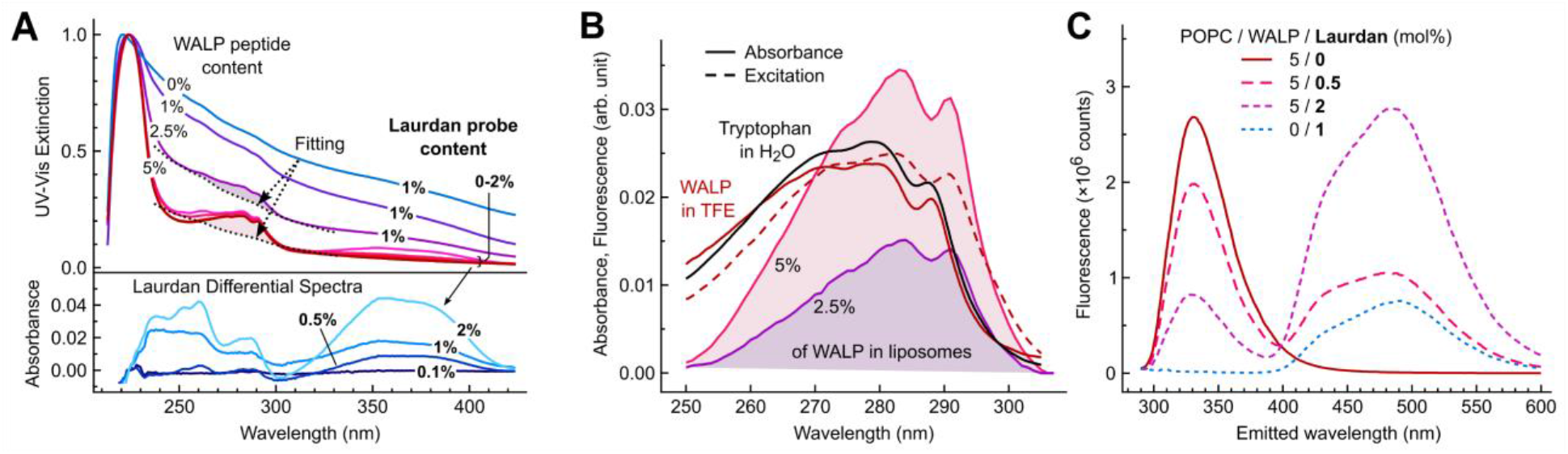
Spectral properties of peptidoliposomes. (A, upper) Extinction spectra measured for different samples. WALP and Laurdan content (mol%) is indicated in the figure. Extinction spectra of liposomes without WALP and Laurdan are indicated by dotted lines. (A, lower) Differential absorbance spectra of Laurdan obtained from peptidoliposomes with 5 mol% WALP and different Laurdan contents (0.1-2 mol%). (B) Absorbance spectra of 2.5 and 5 mol% WALP in liposomes, WALP in TFE, tryptophan in H2O, and excitation spectrum of 5 mol% WALP in liposomes. (C) Fluorescence excitation spectra (at *λ*_EM_=490 nm) of POPC liposomes and peptidoliposomes (5 mol% WALP) with varying Laurdan content (mol%). All spectra were collected at 25°C. *λ*_EX_ and *λ*_EM_ denote excitation and emission wavelengths, respectively

Fluorescence spectra of our samples are shown in Figure 2C. Upon 280 nm excitation, the emission of WALP tryptophan residues appears at 300–390 nm and emission from Laurdan at 400–600 nm. Increasing the Laurdan content in peptidoliposome samples (0, 0.5, and 2 mol%) resulted in an increase in Laurdan emission, and also a decrease in tryptophan emission, clearly suggesting energy transfer between tryptophan and Laurdan. This is evidence of the colocalization of tryptophan and Laurdan and can be used to evaluate membrane association of the peptide, but it does not provide an ultimate proof of peptide incorporation in a transmembrane fashion. It is also important to note, that Laurdan emission was observed upon excitation at 280 nm even in the absence of the peptide, which shows that it does not originate from the energy transfer only (this issue is further discussed in the FRET-GP section).

### GP and TDFS of Laurdan

Laurdan is a well-established solvatochromic probe of lipid membrane.^32, 33^ Its extensive use resulted in variety of the methodological approaches, as well as some inconsistencies in the interpretation of results. Here, we summarize the properties of Laurdan and discuss what it measures in our peptidoliposome samples when excited at its primary excitation band (373 nm).

The light emitted by Laurdan changes its color when the POPC bilayer in which it is embedded is modified (Figure 3A and 3B, sample ➀). A blue-shift is observed upon the addition of 5 mol% of WALP peptide (sample ➁), or to even greater extent upon the addition of 25 mol% of cholesterol (sample ➂). The latter change is partially reversed by heating the sample from 25°C to 37°C (sample ➃). The schematics in Figure 3A illustrates that the color of Laurdan fluorescence reports on lipid packing. Indeed, the response of Laurdan correlates well with the membrane area predicted by computational model.^34^ The two distinct maxima in Laurdan spectra at 440 and 490 nm (Figure 3B) correspond to the emission from rigid and fluid membranes, respectively.

**Figure 3:**
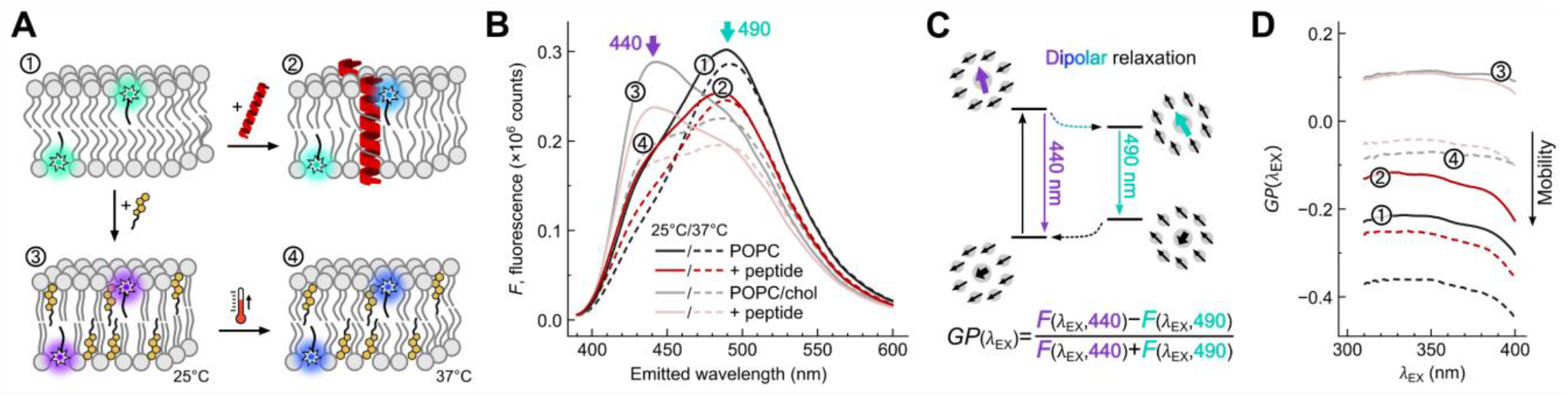
Classical Laurdan GP. (A) Schematic representation of 4 out of the 8 measured systems with symbolic representation of different colors of Laurdan emission. (B) Steady-state emission spectra of Laurdan. Encircled numbers correspond to those of the cartoon in panel A. (C) Scheme of the solvent relaxation process and the resulting time-dependent fluorescence shift as the origin for the distinct Laurdan emission at 440 and 490 nm, and definition of the GP excitation spectra. (D) GP excitation spectra calculated based on Laurdan excitation spectra measured for emission at 440 and 490 nm. Encircled numbers correspond to those of the cartoon in panel A.

Sensitivity to membrane properties arises from the dipolar relaxation (Figure 3C) initiated by the abrupt change in Laurdan dipole moment upon excitation.^35^ The relaxation consists of a rearrangement of Laurdan itself and the hydrated lipids in its vicinity.^36^ The precise location of its fluorophore, slightly below the glycerol backbone of lipids, makes Laurdan directly sensitive to the rearrangement of hydrated lipid carbonyls.^37^ The relaxation process can be comprehensively monitored by measuring Laurdan time-resolved fluorescence, e.g., by recording the entire time-resolved emission spectra and analyzing them using time-dependent fluorescence shift method (TDFS) ^38^. The details of TDFS experiments and their results are discussed in the Supporting Information.

As an alternative to the TDFS approach, one can focus on the intensities of two emitted wavelengths (usually 440 and 490 nm) averaged over time using simple steady-state fluorimeter. The so-called generalized polarization (GP) is a conveniently normalized ratio of these intensities.^28^ It can be calculated as a single number based on the emission spectrum measured for a certain excitation wavelength (e.g., *λ*_EX_ = 373 nm), or it can be calculated for multiple excitation wavelengths at once using two excitation spectra measured for 440 and 490 nm emission, as defined in the equation shown in Figure 3C. In all the samples measured here the addition of the peptide to POPC and POPC/Chol liposomes results in an elevation of the *GP*(*λ*_EX_) spectra (Figure 3D).

Despite attempts to unify the GP method, different interpretations of GP results can be found in the literature.^39^ This interpretations include membrane hydration, structural properties such as lipid packing and ordering, dynamic properties like the mobility of lipids or their parts, and macroscopic parameters like rigidity, viscosity, and fluidity. These properties are often interrelated, for example, an increase in temperature would enhance lipid mobility (dynamics) resulting in membrane expansion (structure), and increased water penetration (hydration). Therefore, it is not always self-evident what GP probes.

Our approach to this problem is based on the comparison of GP results with those of time-resolved measurements (i.e., TDFS), where dynamic properties and hydration are separated.^40^ From our experience the changes in lipid dynamics are the most common. Exclusive changes in lipid hydration are rare and difficult to achieve.^41^ As shown in Figure S1, our GP values represent the rate of dipolar relaxation, rather than hydration. This is a common pattern found in our studies using TDFS,^42^ where we showed that in contrast to peripheral peptide binding, which has little effects on Laurdan, transmembrane peptides reduce lipid mobility without affecting membrane hydration. Here we refer to our GP results, which reflect the mobility of hydrated lipid carbonyls, in more general terms of lipid mobility or membrane fluidity. In this sense, the higher the *GP* value the more rigid the membrane, while low *GP* values represent more fluid membranes. WALP peptide presence rigidifies all of the studied membranes (Figures 3 and S1). This is consistent with our previous study, in which a similar peptide was found to rigidify lipid membranes and reduce lipid diffusivity.^29^ In that study the effect of the peptide was enhanced by cholesterol, which segregated from the peptide surface. In the present study, the influence of cholesterol is within the experimental error, possibly due to different lipid tails used.

In is important to consider the limitations of Laurdan sensitivity when interpreting these results. For example, in rigid membranes, the relaxation times become longer than the fluorescence lifetime of Laurdan, diminishing the changes GP values. This is evident in Figure 3D, where the least fluid samples are those containing cholesterol and measured at 25°C. In this case, the addition of WALP does not change *GP* significantly. For the most fluid systems (no cholesterol, 37°C) addition of WALP rigidifies the membrane markedly. Contrary to our findings, Dinic et al. conclude from their measurements performed for two transmembrane peptides that Laurdan was not sensitive to their presence.^43^ However, a closer look at the emission spectra presented suggests that some rigidification may have been present in their system as well.

### Energy Transfer

Laurdan emission upon tryptophan excitation (*λ*_EX_ = 280 nm) suggested that Laurdan acted as an acceptor of Förster resonance energy transfer (FRET) from the WALP peptide (Figure 2C). The overlap between WALP tryptophan emission and Laurdan excitation is indeed substantial (Figure 4A), but the efficiency of FRET diminishes with the sixth power of the distance between the molecules.^44^ The distance above which FRET efficiency drops below 50% (Förster radius, *R*_0_) is 2.7 nm (see the Supporting Information for details), which is in good agreement with the previously reported values for Laurdan in egg PC vesicles with octyl-tryptophan (3.1 ± 0.1 nm) and the nicotinic acetylcholine receptor (2.9 nm).^27^ Assuming a typical alpha helix diameter^45^ of 1.2 nm and the distance of two adjacent POPC lipids of 0.9 nm (based on an area per lipid of 0.7 nm^2^ and hexagonal lipid packing),^46^ our *R*_0_ covers two shells of lipid molecules around the peptide and reaches the third, i.e. (2.7 − 1.2/2)/0.9 = 2.3 diameters of the POPC molecule. This is schematically depicted in Figure 5C, in which the *R*_0_ range is marked as red dashed circles. The distance between the two tryptophan pairs in a WALP molecule – 2.85 nm (19 AA × 0.15 nm) matches very well the distance of 2.67 nm between the carbonyls of the two opposing leaflets of POPC bilayer that are probed by Laurdan,^46^ especially considering possible slight tilts of the peptide from the membrane normal. This means that the depths of Laurdan and WALP tryptophan moieties location in the POPC membrane coincide. Also, since the distance of the tryptophan pair on one side of the lipid bilayer to the Laurdan on the other side is approximately equal to *R*_0_, FRET between the two bilayer leaflets is probable.

**Figure 4:**
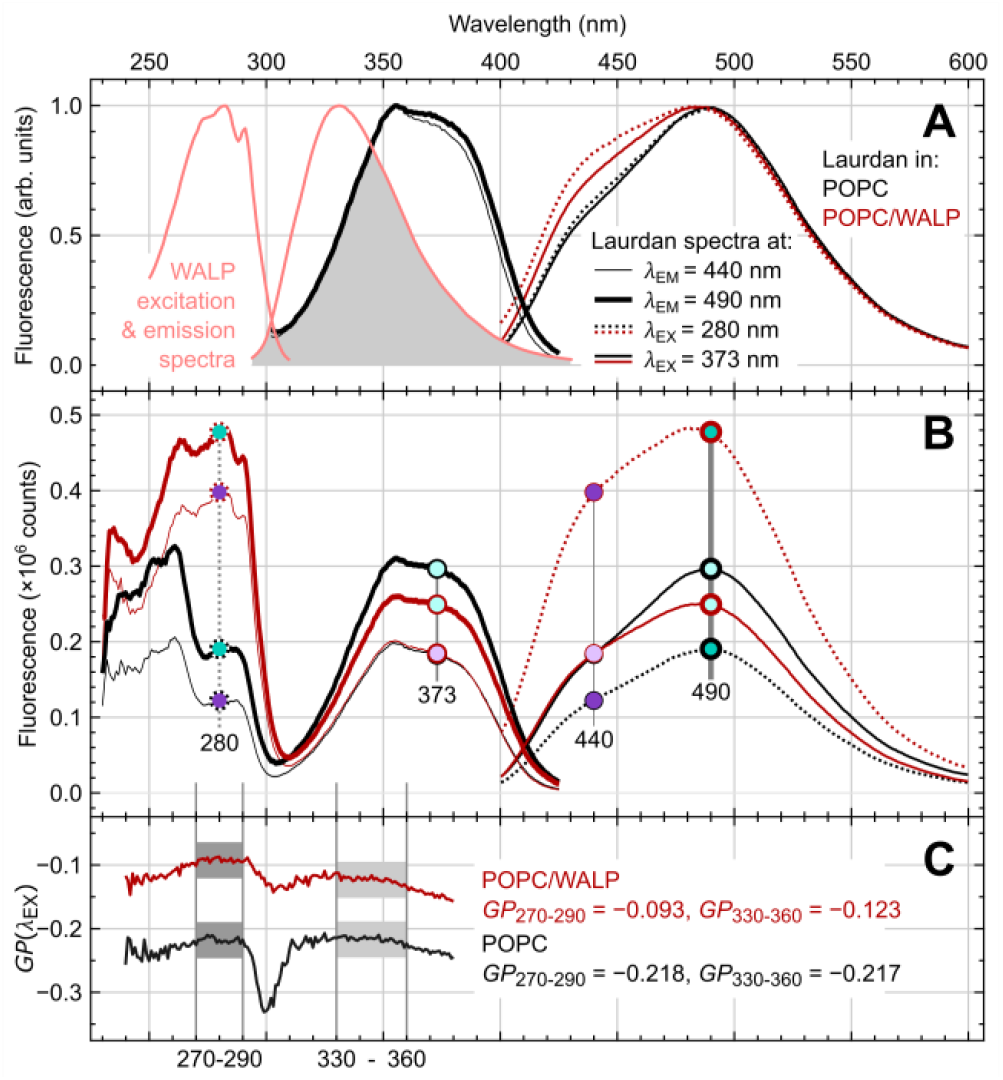
Laurdan GP initiated by FRET from tryptophan. (A) Normalized excitation and emission fluorescence spectra of WALP peptide and Laurdan embedded in POPC vesicles. Spectral overlap between WALP emission and Laurdan excitation marked in gray. Variability of the shape of Laurdan spectra is depicted for different excitation (*λ*_EX_) and emission (*λ*_EM_) wavelengths. (B) Full-range excitation spectra (for *λ*_EM_ = 440 and 490 nm) and emission spectra (for *λ*_EX_ = 280 and 373 nm) of Laurdan in POPC and in POPC/WALP membranes. Fluorescence intensities used to calculate GP values are marked both in the excitation and in the emission spectra; same fill colors are used for the corresponding points in both spectra. (C) Full-range GP spectra calculated from the excitation spectra shown in panel B. The values of *GP*_270-290_ and *GP*_330 360_ are given. The regions of GP spectra from which they were obtained are marked in gray.

**Figure 5:**
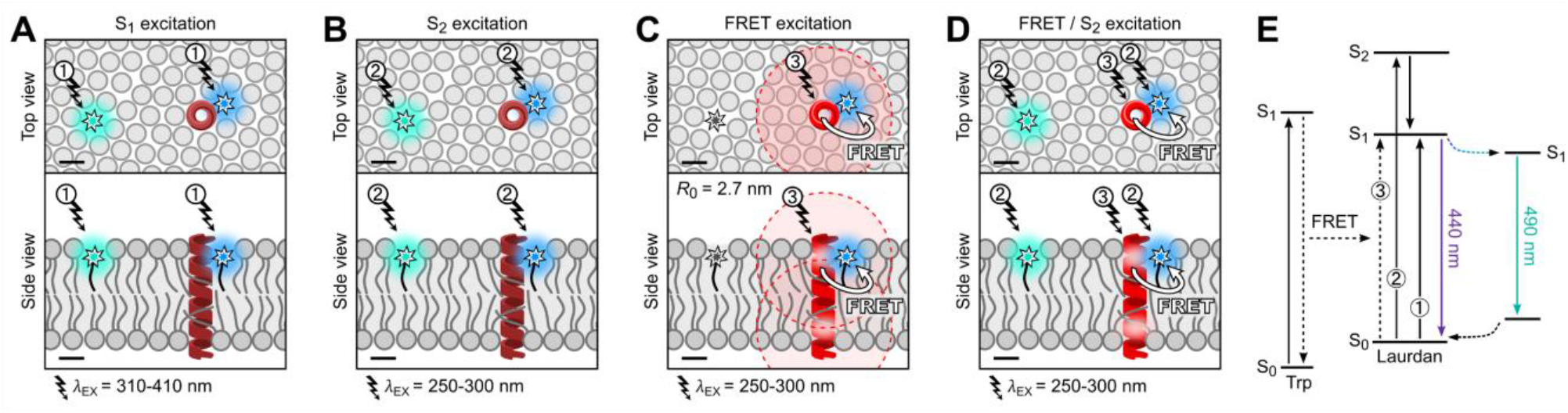
Different scenarios of Laurdan excitation in peptidoliposomes. (A) Standard S_1_ excitation. (B) S2 excitation. (C) FRET from peptide tryptophan to Laurdan. (D) Concurrent FRET and S_2_ excitation of Laurdan. (B) Jablonski diagram of different pathways of Laurdan excitation and the subsequent dipolar relaxation. Encircled numbers correspond to different excitation paths: ➀ Direct Laurdan excitation to the S_1_ state, ➁ Direct Laurdan excitation to the S_2_ state, ➂ Excitation of Laurdan via FRET from peptide tryptophan (Trp). Laurdan fluorophores are depicted as stars, WALP peptide as a red helix, and excitation light as bolt arrows. Förster radii of 2.7 nm are indicated with dashed circles around the tryptophan residues. Scalebars in the bottom left corners are 1 nm.

Bearing in mind the precise location of Laurdan in the membrane, the observed FRET indicates that the peptide or protein must be either closely adsorbed to the membrane or embedded in it. Knowing *R*_0_, and measuring FRET efficiency one can predict the distance between donor and acceptor molecules in a sample of interest. An additional requirement for this prediction is an assumption of certain distribution of these molecules. A robust way of obtaining this for membranes is to use Monte Carlo simulations and compare the calculated and the measured fluorescence decays.^47-49^ As a simple alternative we assume a random distribution of donors and acceptors in the membrane and estimate an average distance between them as:

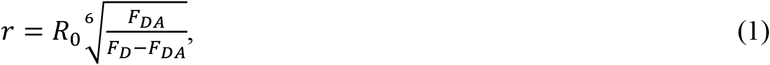

where *F*_D_ and *F*_DA_ are fluorescence intensities of the donor in the absence and the presence of the acceptor, respectively. FRET efficiency and the corresponding *r* values calculated using Eq. 1 are depicted in Table 1.

**Table 1:** Tryptophan-Laurdan FRET efficiency and donor-acceptor distances.

| Laurdan content (mol%) | FRET efficiency (%) | Donor-acceptor distance, $r^a$ (nm) |
| --- | --- | --- |
| 0.1 | 11 | 3.9 |
| 0.2 | 14 | 3.7 |
| 0.5 | 26.5 | 3.2 |
| 1 | 40.5 | 2.9 |
| 2 | 69 | 2.4 |

<sup>a</sup> calculated according to Eq. 1.

### FRET-GP – Origin

The excitation of Laurdan via FRET preserves its fluorescence properties and its sensitivity to lipid hydration and mobility. This is evident from the variability of the Laurdan emission spectra measured for POPC and POPC/WALP samples at 25°C (Figure 4A). The emission spectra of Laurdan excited in peptidoliposomes at 280 nm can be analyzed in the same way as those recorded for the classical excitation. More interestingly, the GP results obtained this way (*GP*_280_) differ from the classical ones (*GP*_373_). For POPC, the emission spectra at these two excitation wavelengths are similar, but they diverge significantly for POPC/WALP. The resulting GP values were: *GP*_373_ = −0.24, *GP*_280_ = −0.22 for POPC, and *GP*_373_ = −0.15, *GP*_280_ = −0.09 for POPC/WALP. In other words, the presence of WALP elevates both *GP*_373_ and *GP*_280_, but the magnitude of the change is considerably higher for *GP*_280_. Qualitatively comparable results were obtained for the nicotinic acetylcholine receptor in egg PC vesicles for similar excitation wavelengths (360 nm for direct and 290 nm for FRET condition)^.27^

Measuring GP spectra (as in Figure 3D) is a more robust way of analyzing Laurdan fluorescence than calculating a single *GP* value from an emission spectrum.^50^ In our case, the GP spectrum has the additional advantage of showing, in a single curve, the GP values for both the classical Laurdan excitation and the wavelengths typical for tryptophan excitation. For POPC and POPC/WALP, the full-range fluorescence excitation spectra measured from 200–430 nm for the emission at 440 and 490 nm are depicted in Figure 4B together with the corresponding emission spectra (the same as those shown in Figure 4A, but before intensity normalization). Fluorescence intensities for the characteristic wavelengths (*λ*_EX_ = 280 and 373 nm, *λ*_EM_ = 440 and 490 nm) are marked in both excitation and emission spectra. Full-range GP spectra calculated from the excitation spectra from Figure 4B are shown in Figure 4C. When analyzing such GP spectra, we choose to average the flat parts corresponding to two excitation peaks, i.e., 270–290 nm and 330–360 nm, to get *GP*_270-290_ and *GP*_330-360_, respectively. The latter is the standard measure of GP, while the first is expected to provide the GP arising from the FRET from tryptophan. These values correspond well to the *GP*_373_ and *GP*_280_ mentioned above, but as the averaged values are less prone to deviations. Before describing the GP results, we need to address an important issue, namely, the origin of Laurdan excitation at 280 nm in the absence of any peptide. Clearly, this excitation cannot originate from FRET in the absence of the donor. Instead, the excitation spectrum of Laurdan between 230 and 300 nm corresponds to the direct excitation of the probe to its second singlet state (S_2_). As we discuss below, S_2_ excitation of Laurdan contributes to the measured *GP*_280_ and *GP*_270-290_, and should be always considered when analyzing FRET-GP results.

### Different Scenarios of Laurdan Excitation

When analyzing fluorescence of Laurdan in the presence of tryptophan, the following scenarios should be considered. Firstly, direct excitation of Laurdan to its first singlet excited state (S_1_) is achieved by UV to deep violet light (310–410 nm). This excitation is marked as ➀ in Figure 5A and the Jabłoński diagram in Figure 5E. The dipolar relaxation that follows gradually lowers the energy of Laurdan together with its nanoenvironment from S_1_ to S_1_’, which can be monitored by the emitted fluorescence (violet to green). As illustrated in Figure 5A, Laurdan nanoenvironments may differ, but the response from the whole sample is uniformly averaged.

The second excitation scenario (Figure 5B) is similar to the first one, but is initiated by UV in the range of 250– 300 nm, which causes S_0_ → S_2_ transition of Laurdan; marked as ➁. Since the lifetime of the S_2_ state is short (typically in the ps range) it quickly relaxes to the S_1_ state^44^ and the dipolar relaxation advances in an almost identical way to that initiated by S_1_ excitation. This is why, in the absence of tryptophan, *GP*_S1_ ≈ *GP*_S2_, or at the measurement conditions *GP*_270-290_ ≈ *GP*_330-360_.

The third scenario requires the presence of tryptophan, which can be excited by 250–300 nm UV to its S_1_ state and subsequently donate its energy to Laurdan S_0_ → S_1_ transition; marked ➂. As illustrated in Figure 5C, only the Laurdan molecules located within the Förster radius of the peptide can be excited this way. Such Laurdan excitation is delayed relative to the tryptophan excitation by the time that tryptophan spends in its S_1_ state before the FRET occurs. Nevertheless, once excited, Laurdan reaches its S_1_ state and the process continues as in the first scenario. In other words, the FRET process does not affect the relaxation monitored by GP measurements. This enables Laurdan to selectively monitor peptide nanoenvironments.

Unfortunately, the second and the third scenarios happen at the same excitation range (Figure 5D), therefore *GP*_270-290_ measured for our peptidoliposomes contains both FRET and S_2_ contributions; we refer to them as *GP*_FRET_ and *GP*_S2_. If, like in our case, the peptide rigidifies the membrane locally more than globally (*GP*_FRET_ > *GP*), the *GP*_S2_ contribution weakens the effect measured as *GP*_270-290_. This means that *GP*_FRET_ should be even greater than what we measure as *GP*_270-290_. The same is true for the results obtained in the work by Antollini et al.^27^ who measured GP at 290 nm excitation disregarding S_2_ contribution. The S_2_ contribution is inevitable and was also present in that work as evident from figures 3a and 8,^27^ which include Laurdan emission spectra measured at 290 nm excitation in the absence of tryptophan. To obtain *GP*_FRET_, one should remove the S_2_ contribution from the measured *GP*_270-290_.

### FRET-GP – Removal of the S_2_ Contribution

The separation of *GP*_FRET_ and *GP*_S2_ contributions from *GP*_270-290_ is a challenging task. Subtracting blank measured for a peptide-free sample is not a viable solution, since the peptide changes membrane fluidity characterized by *GP*_S2_. A more accurate approach would be to replace tryptophan residues with different amino acids to create a control sample. However, this approach requires additional measurements and considerable effort in peptide/protein engineering – design, production and testing of a new peptide/protein.

FRET and S_2_ contributions can be separated by spectral demixing, if only they produce excitation spectra of Laurdan that sufficiently differ from each other. Figure 4B shows that the shapes of the excitation spectra for liposomes and peptidoliposomes are indeed distinct in the range 250–300 nm. The spectra for liposomes represent S_2_ excitation only, while those for peptidoliposomes represent the sum of S_2_ and FRET contributions. When analyzing S_2_ excitation spectra measured for different POPC-based lipid systems in the absence of peptide, we found that their shape is largely conserved, with different systems differing mostly in their amplitudes. Interestingly, the choice of temperature (i.e., 25°C and 37°C) or emission wavelength (440 or 490 nm) also did not significantly influence the shape. This provides an opportunity to determine a universal lineshape for Laurdan S_2_ excitation, *L*_S2_(*λ*_EX_). Among the tested systems, the shapes of the excitation spectra measured for the samples containing cholesterol were the most distinct. Therefore, we decided to process them separately. Limiting our analysis to 245-300 nm and normalizing the intensity of all excitation spectra to 0.5 on average, we obtained *L*_S2_(*λ*_EX_) separately for cholesterol-free and cholesterol-containing samples (black and gray curves in Figure 6A, respectively). After subtracting these lineshapes from the excitation spectra for WALP peptidoliposomes normalized to 1, we obtained FRET excitation lineshapes, *L*_FRET_(*λ*_EX_), separately in cholesterol-free and cholesterol-containing samples (dark and light red curves in Figure 6A, respectively). The lineshapes were smoothed in OriginPro 2021b using a Savitzky-Golay filter with a 7-point window. They are provided in Appendix S1. The S_2_ lineshapes appear slightly different depending on the presence of cholesterol in the system, while FRET lineshapes do not. FRET lineshapes are also very similar in shape to the excitation spectrum of WALP measured at the emission wavelength of tryptophan (330 nm; dashed line in Figure 6A). It is important to note that the lineshapes may differ for different peptides/proteins and for different lipid systems (see, e.g., Figure S4A).

**Figure 6:**
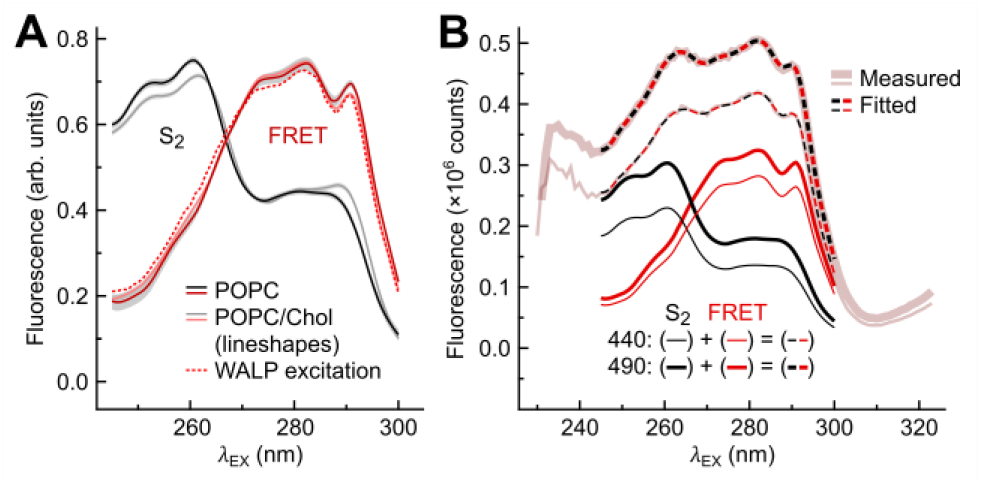
Removal of Laurdan S_2_ contribution by spectral demixing. (A) Laurdan S_2_ and FRET excitation spectral lineshapes for POPC and POPC/cholesterol membranes. Excitation spectrum of WALP tryptophans measured at *λ*_EM_ = 330 nm emission is shown as dotted line for comparison. All spectra are normalized to 0.5 on average. Gray areas represent standard error. The lineshapes are also reproduced in Appendix S1. Their comparison with the lineshapes obtained for DMPC at different temperatures can be found in Figure S4A. (B) Spectral demixing from the measured excitation spectra. Fitting of the linear combination of S_2_ (black lines) and FRET (red lines) lineshapes from panel A to the measured excitation spectra of Laurdan in POPC/WALP membranes separately for *λ*_EM_ = 440 and *λ*_EM_ = 490 nm (see the main text for details).

Having determined *L*_S2_(*λ*_EX_) and *L*_FRET_(*λ*_EX_) lineshapes, we fitted their linear combinations to the measured excitation spectra:

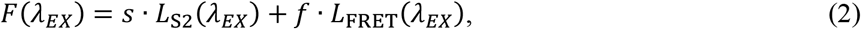

where *s* and *f* are non-negative constants obtained from the fitting. The fitting is performed separately for the excitation spectra measured at 440 nm and 490 nm emission, giving *s*_440_, *f*_440_, *s*_490_, and *f*_490_ values. From these parameters, *GP*_S2_ and *GP*_FRET_ can be calculated:

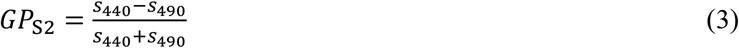

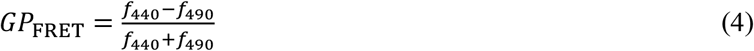

As exemplified in Figure 6B depicting this procedure, the fitted curves completely overlap with the measured data. Having *s*_440_, *f*_440_, *s*_490_, and *f*_490_ one can also easily calculate the percentage of FRET contribution to Laurdan emission:

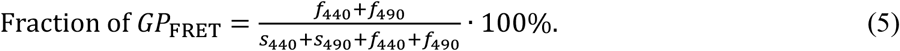

Based on the above-presented approach, we also proposed a simplified analytical method for *GP*_FRET_ determination based on fluorescence intensities at individual wavelengths. Derivation of this method and its validation is provided in the Supporting Information.

*GP*_FRET_ exclusively characterizes the membrane regions close to a peptide or protein. The interpretation of GP_S2_ is, however, more complicated. In principle, all of the Laurdan molecules can be excited to their S_2_ state and contribute to *GP*_S2_. This means that *GP*_S2_ can give values close to classical GP representing the whole membrane in a more or less uniform fashion. Only in the systems where a substantial fraction of all Laurdan molecules is excitable via FRET, *GP*_S2_ would overrepresent the regions far from the donor. It is because close to the donor Laurdan molecules that are being excited via FRET are temporarily unavailable for the S2 excitation, which lowers their contribution to *GP*_S2_. Thus, selectivity of *GP*_S2_ depends on the concentration of donors and Laurdan as well as their distribution in the membrane.

Overall, the removal of the S2 contribution proposed here is a straightforward procedure. It requires only Laurdan excitation spectrum measured in the absence of donor to obtain the *L*_S2_(*λ*_EX_) lineshape. In case such measurement is not feasible, one can first determine *L*_FRET_(*λ*_EX_), assuming it to be of the shape of the peptide/protein emission spectrum (*λ*_EX_≈330 nm), and then subtract *L*_FRET_(*λ*_EX_) from Laurdan excitation spectrum measured in the system of interest.

### FRET-GP – Results and Their Meaning

*GP*_FRET_ and *GP*_S2_ obtained from Eq. 3 and 4 after fitting appropriate lineshapes to the measured excitation spectra are shown in Figure 7 together with classical GP (*GP*_330-360_) and the uncorrected FRET-GP (*GP*_270-290_) for comparison. In general, the presence of peptide increases all *GP* parameters. *GP*_330-360_ increases with WALP content (0, 1, 2.5, and 5 mol%) to −0.217, −0.192, −0.174, and −0.123, respectively. For the wavelength region in which FRET-GP occurs (*GP*_270-290_) the values are higher, except for the sample without peptide. Additionally, *GP*_270-290_ increases faster than *GP*_330-360_ with peptide content, being −0.001, 0.016, 0.024, and 0.038 larger for 0, 1, 2.5, and 5 mol% peptide, respectively. These data contain contributions from both FRET-GP and GP from the S_2_ state. However, they already provide valuable additional information. Not only is it clear that the peptide reduces the fluidity of the POPC lipid bilayer, but also that the effect is more pronounced locally.

**Figure 7:**
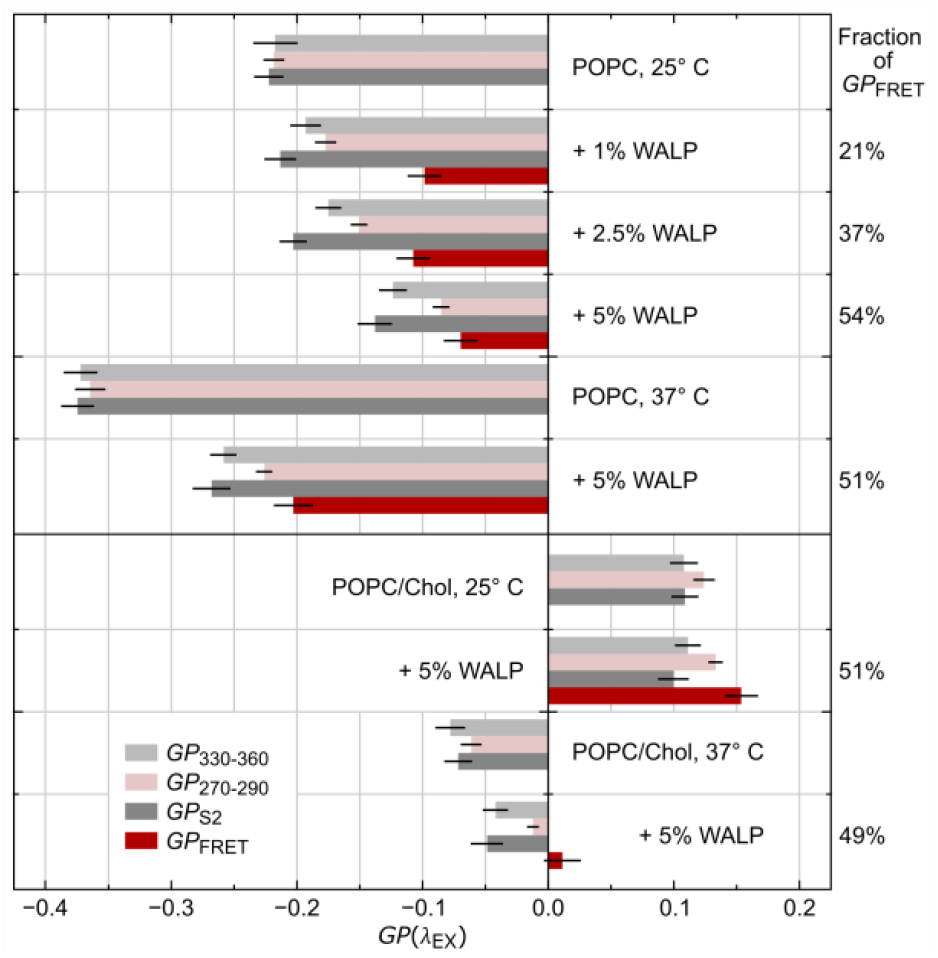
Laurdan GP and FRET-GP results for POPC-based vesicles. *GP* values obtained for different WALP content in POPC and POPC/cholesterol membranes measured at 25°C and 37°C. *GP*_270-290_ and *GP*_330-360_ were obtained directly from the full-range GP spectra, while *GP*_FRET_ and *GP*_S2_ from the procedure of spectral demixing. Percentage of FRET contribution to Laurdan fluorescence intensity is given on the right side of the figure. Error bars represent standard deviation.

Removal of the S_2_ contribution significantly enhances this difference between *GP*_FRET_ and *GP*_330-360_. Already 1 mol% WALP added to POPC causes its increased from the initial *GP*_330-360_ = −0.217 to *GP*_FRET_ = −0.123, compared to *GP*_270-290_ = −0.201. *GP*_FRET_ is 0.094, 0.067, and 0.053 above *GP*_330-360_, for 1, 2.5, and 5 mol% peptide, respectively. It is important to note that *GP*_270-290_ can be measured for any Laurdan sample, while *GP*_FRET_ cannot be determined for samples that do not contain peptide. *GP*_FRET_, as a local measure, is much less dependent on the peptide content than *GP*_330-360_. These results highlight the importance of the S_2_ removal. In the previous study^27^ ∼50 wt% protein enrichment^51^ of egg PC produced ∼0.06 increase in GP when measured at FRET conditions, which is half of the effect we observe in *GP*_FRET_ for 1 mol% WALP peptide (∼5 wt%) at the same temperature.

*GP*_S2_ is 0.005, 0.021, 0.028, and 0.015 lower than *GP*_330-360_ for 0, 1, 2.5, and 5 mol% peptide, respectively. This means that membrane fluidity further apart from the peptide is higher than the average. The difference becomes initially more pronounced with peptide content, when the FRET contribution increases. But for the highest peptide content it decreases again, since in this case many Laurdan molecules are inevitably close to the peptides.

As shown on the right of Figure 7 the contribution of FRET to the total Laurdan fluorescence intensity increases gradually with the peptide content, but saturates at the highest peptide content. This is because, even in the case where FRET efficiency is 100% and all Laurdan molecules are available for FRET, some of them will still be excited directly to the S_2_ state.

All the results discussed above were measured at 25°C. At 37°C, the membranes are more fluid, which results in lower GP values. However, the presence of the peptide rigidifies the membrane in a similar manner at both temperatures. The rigidification caused by the peptide at 37°C is comparable to that caused by cooling POPC membrane by 12 degrees, namely *GP*_FRET_ for POPC/WALP at 37°C is at the level of GP for pure POPC at 25°C.

In cholesterol-containing systems, all GP values are elevated, indicating that the membrane fluidity in general, and the lipid carbonyl mobility in particular, are reduced. Even in these systems, the addition of peptide alters the GP values. At 25°C, the change in standard GP is within the error, and that of *GP*_270-290_ is at the border of the error. However, the increase in *GP*_FRET_ is statistically significant. At 37°C, all of the GP values are elevated by the peptide, but the most pronounced change is again in *GP*_FRET_.

### FRET-GP – Lipid Phase Transition in Peptidoliposomes

We further tested FRET-GP approach by probing the thermotropic phase transition of DMPC in peptidoliposomes with 5 mol% of WALP peptide. Pure DMPC liposomes were used as a reference. The results are presented in Figure 8. Upon heating, the emission of Laurdan gradually red-shifts and lowers all presented GP parameters, but their profiles are distinct. Laurdan emission spectra obtained by excitation with 280 nm red-shift to larger extent when the peptide is present (Figure 8A and 8B). The main P_β_-L_α_ phase transition of DMPC is clearly visible in *GP*_330-360_ curve and its first derivative (Figure 8C and 8D, respectively). Its temperature determined here as T_m_ = (24.0 ± 0.5)°C agrees with the literature value T_m_ = (23.6 ± 1.5)°C. WALP presence makes the phase transition less sharp, but the first transition still allows to identify the position of the inflection point as T_m_ = (23.2 ± 0.5)°C. Reduction of T_m_ of DMPC was previously measured for similar transmembrane peptide using EPR.^52^ *GP*_FRET_ reports even slightly more reduced T_m_ at the peptide, i.e., (22.9 ± 0.5)°C. Similarly to POPC/WALP, *GP*_FRET_ > *GP*_330-360_ for DMPC/WALP. This is true not only for the liquid crystalline, but also for the gel phase. Close to T_m_, all the GP parameters are indistinguishable for pure DMPC, likely due to membrane defects expected at the phase transition. Overall, WALP rigidifies the DMPC membrane, stabilizes lipid membrane properties against temperature changes and slightly lowers the main phase transition temperature.

**Figure 8:**
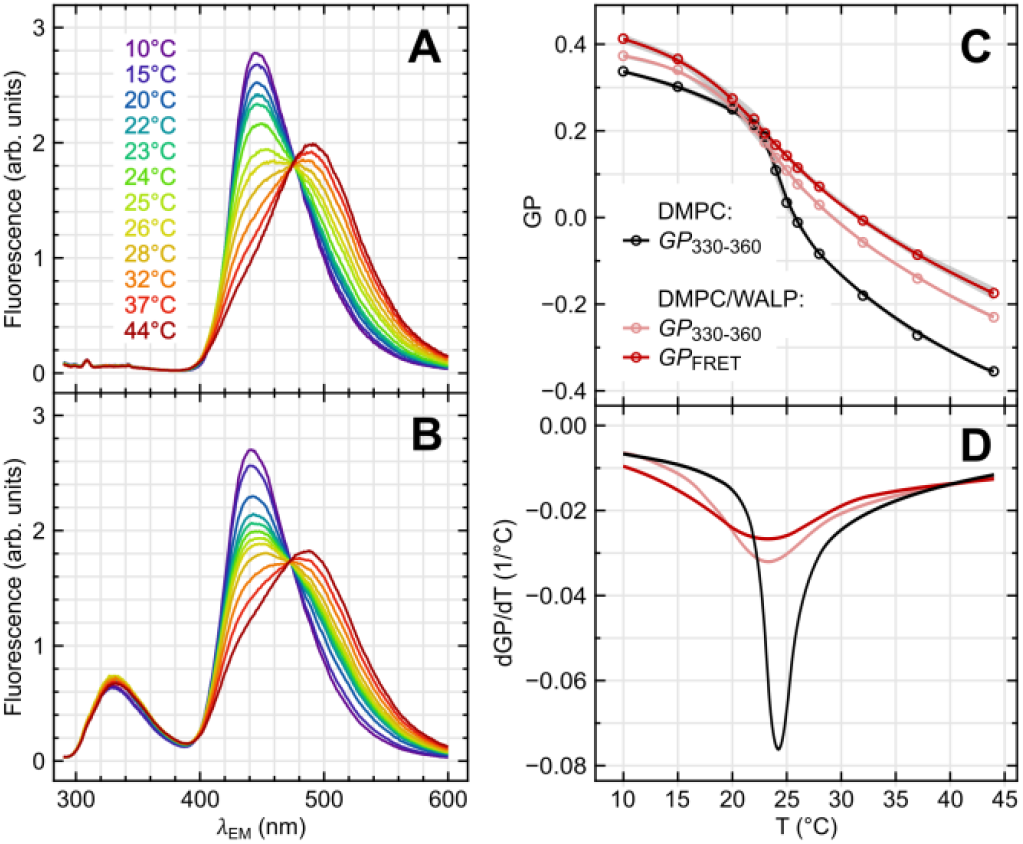
Laurdan GP and FRET-GP results for DMPC-based vesicles at different temperatures. Emission spectra at *λ*_EE_ = 280 nm of Laurdan embedded in DMPC liposomes (A) and DMPC/WALP peptidoliposomes (B). Spectra were normalized for the area between 400 and 600 nm. (C) Dependence of classical GP_330-360_ and GP_FRET_ on the temperature. Gray areas represent standard deviation. (D) First derivative of the GP curves from panel (C).

Laurdan S2 lineshapes used for spectral demixing change considerably for DMPC gel phase. This motivated us to address the lineshape issue in more details (see the Supporting Information). In short, while the quality of the lineshape fit performed with POPC-derived lineshapes was markedly low for the gel phase samples, the obtained *GP*_FRET_ results were still in perfect agreement with those obtained using proper DMPC-derived lineshapes.

### FRET-GP – Limitations and Outlook

There are a few limitations that should be considered when planning FRET-GP experiments. First, the method requires the presence of tryptophan residues in the peptide or protein of interest. However, the existence of multiple tryptophan and tyrosine residues can sometimes obscure the excitation spectrum of a protein. This could, in turn, impede the removal of the S_2_ contribution, and consequently reduce the precision or specificity of the obtained *GP*_FRET_ values. Second, the low Förster radius for Laurdan and tryptophan increases the precision of the peptide vicinity selection, but reduces FRET efficiency and demands appropriate tryptophan location. Favorably, in many transmembrane proteins, tryptophan is present in the so-called aromatic belt,^13, 53^ which is located at the same depth as the fluorophore of Laurdan.^37^ This highly increases the prospect of sufficient FRET efficiency.

Applicability of FRET-GP is not limited to liposomes. The unique sensitivity and specificity of the method allows investigation of many aspects of protein-lipid interactions in various model lipid systems, including nanodiscs and bicelles. Labelling of the lipid bilayer with a small solvatochromic probe while keeping the protein structure intact minimizes system disturbance.

FRET-GP can be easily extended to different FRET pairs. Many other fluorescent polarity probes besides Laurdan possess excitation properties suitable for accepting FRET from tryptophan, e.g., Patman, C-Laurdan, and Prodan. The different characteristics of these probes such as, charge, gives opportunity to control the affinity of the probe to a protein of interest. Additionally, since different polarity probes are located at different depths it the bilayer, it should be possible to specifically monitor the influence of a transmembrane domain in different regions of the lipid bilayer, e.g., separately in the hydrophobic core and in the headgroup region.

It is possible to specifically label a protein of interest with a fluorescent marker excitable with visible light and use another long-wavelength fluorescent probe of polarity. This approach would be beneficial for live-cell studies and microscopy, which are difficult for the tryptophan-Laurdan pair. The first difficulty is the abundance of proteins in living systems, which would impair the specificity of the method in the current form. Secondly, porting the method to fluorescence microscopy requires either a sophisticated UV microscope or multiphoton excitation of tryptophan. Popular Ti-sapphire lasers are capable of 3-photon excitation of tryptophan. In this case, however, a further obstacle is the fact that the same wavelength that is used to 3-photon excite tryptophan, would also be able to 2-photon excite Laurdan. This issue might be also an overlooked source of artifacts in Laurdan microscopy. When 2-photon excitation of Laurdan in living cells is used, one might also unintentionally excite tryptophan in membrane proteins with the same wavelength (via 3-photon excitation). In such cases, the measured GP would contain a FRET-GP contribution. All of these issues require additional studies to be properly addressed.

### Are These Annular Lipids What FRET-GP Probes?

Our results show that WALP significantly decreases membrane fluidity, which is common effect for transmembrane peptides.^42^ The molecular mechanism responsible for this is the trapping of the lipid acyl chains at the rough surface of the transmembrane domain, as proposed previously ^29^. As a consequence, the lipid mobility probed by Laurdan, as well as the lateral diffusivity of lipids is reduced, while the order of lipid tails increases.

These findings were based on measuring average membrane properties. The present study advances our understanding of lipid interactions with transmembrane domains. The effects measured using FRET-GP are stronger, and thus, noticeable at lower peptide concentrations. More importantly, the specificity of probing lipids in the immediate surrounding of the peptide reveals lateral inhomogeneity in lipid fluidity and sheds new light on the concept of annular lipids. The concept originated soon after the introduction of the influential Singer-Nicolson model defining protein embedding in the lipid bilayer,^54^ when hindrance of lipid mobility was detected using EPR.^55^ The idea that lipids form a ring (or annulus) surrounding transmembrane domain of a protein that has different properties from the bulk lipids was coined as an analogy to solvation shell. While still widely used today^8^ it is also strongly criticized by some.^56^ In this respect, FRET-GP confirms that the mobility of lipids within 2-3 layers from the transmembrane peptide is significantly reduced compared to the bulk lipids. The consequences of these local effects are, however, global – the average mobility of lipids in the membrane is reduced. FRET-GP also proves that even the lipids that are in immediate contact with the peptide are still mobile.

The experiments with the phase transition in DMPC/WALP membrane revealed that the peptide acts as a buffer against temperature changes, stabilizes membrane fluidity, and slightly lowers the phase transition temperature. Overall, our findings support the view of lipid annulus as a solvation shell of membrane protein, where the lipids are slowed down by the transmembrane domain, but still mobile and readily exchangeable with the bulk lipids.

## CONCLUSIONS

When exciting Laurdan via FRET from tryptophan an unwanted contribution from direct Laurdan excitation to its S_2_ state inevitably contaminates the GP results. We propose a simple procedure that alleviates this issue and allows FRET-GP to specifically determine the local and global impact of a transmembrane protein or peptide on lipid membrane fluidity. The method combines the spatial precision of the nm range of FRET from tryptophan – the natural endogenous fluorophore – and the unsurpassed sensitivity of Laurdan to the changes in lipid mobility. Tryptophan-Laurdan combination is also optimal due to its good spectral match and the location of the fluorophore of Laurdan exactly at the typical position of the aromatic belt of many transmembrane proteins. Considering the importance of membrane proteins and their interactions with lipids for biology and pharmaceutical research, FRET-GP as a fast and reliable method for studying them can be of broad interest.

Our results demonstrate that WALP reduces the fluidity all lipids, especially those in the first two shells surrounding the peptide. Additionally, the temperature of the main lipid phase transition of DMPC was broadened and lowered by the peptide. These findings pair well with the mechanism of lipid acyl chain entrapment at the rough surface of a protein transmembrane domain.

## ASSOCIATED CONTENT

### Supporting Information

DLS – additional results and discussion, TDFS – method and results, Förster radius calculation method, simplified point method of S2 removal – method and results, S2 and FRET lineshapes for DMPC – additional results and discussion, POPC and POPC/Chol lineshapes.

## AUTHORINFORMATION

### Author Contributions

G. C. N. T. and A. U. performed the experiments, analyzed the data, contributed to figure preparation and manuscript writing and editing. P. J. conceived the study, designed data analysis, contributed to data processing, prepared the figures and wrote the manuscript. G. C. N. T. and A. U. contributed equally to this work.

## ACKNOWLEDGMENTS

We thank Federica Scollo for critical reading of the manuscript and many valuable comments, and Marek Cebecauer, Martin Hof, and Radek Sachl for fruitful discussions. We acknowledge financial support from the Czech Science Foundation (EXPRO grant 19-26854X).

## ABBREVIATIONS

Chol: cholesterol
DLS: dynamic light scattering
DMPC: dimyristoylphosphatidylcholine
EDTA: ethylenediaminetetraacetic acid
egg PC: phosphatidylcholine (egg, chicken)
EPR: electron paramagnetic resonance
FRET: Förster resonance energy transfer
FRET-GP: Förster resonance energy transfer-generalized polarization
GP: generalized polarization
HEPES: 4-hydroxyethyl piperazineethanesulfonic acid
IR: infrared
Laurdan: 6-lauroyl-2-dimethylaminonaphthalene
NMR: nuclear magnetic resonance
POPC: palmitoyloleoylphosphatidylcholine
TCSPC: time-correlated single photon counting
TDFS: Time-dependent fluorescence shift
TFE: trifluoroethanol
WALP: GTSTSKKWWLALALALALALALALALWWKKFSTS peptide.

