## Supplementary Information for "FRET-GP – A Local Measure of the Impact of Transmembrane Peptide on Lipids"

### TABLE OF CONTENTS

1. Dynamic light scattering (additional results and discussion)
2. Time-dependent fluorescence shift (method and results)
3. Förster radius calculation (method)
4. Simplified point method for the S2 contribution removal (method and results)
5. $S_2$ and FRET lineshapes for DMPC phase transition (additional results and discussion)
6. References
7. Appendix – POPC and POPC/Chol lineshapes

### 1. DYNAMIC LIGHT SCATTERING

When characterizing liposomal formulations of calibrated size dynamic light scattering (DLS) allows to easily spot many problems, e.g., the presence of cholesterol crystals precipitated from a lipid bilayer. The size distribution of sonicated liposomes is usually broader than that of extruded ones, which limits the precision of DLS characterization. The size distribution results from the fitting of a model to the autocorrelation function calculated from the recorded light scattering. For the samples with varying particle sizes, the models are often not fully applicable, which can considerably distort the results. Therefore, we often use simplified cumulant analysis for sonicated proteoliposomes, and always examine the goodness of the fit and the shape of the original autocorrelation functions. The most common defects of proteoliposome morphology can be observed directly in the autocorrelation curves prior to any fitting. The outliers of the expected vesicle sizes cause deviations in the beginning or in the tail of the autocorrelation function. These two parts of the curve are usually flat and featureless, therefore any deviation can be easily spotted. Our data show no signs of peptide precipitation nor vesicle aggregation, except in the POPC/1 mol% WALP sample, for which a long and rough autocorrelation tail indicates substantial vesicle aggregation, and results in large vesicle sizes with large standard deviation (Figure 1B).

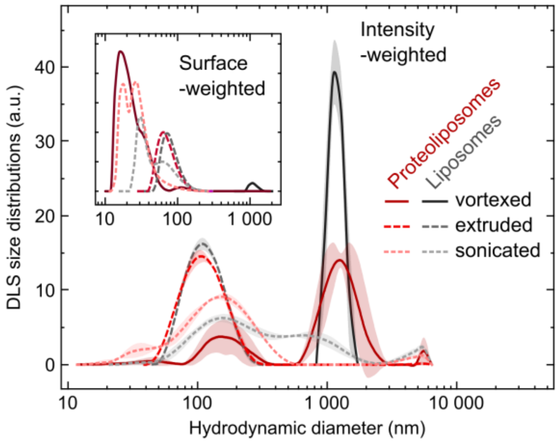


**Figure S1**. Intensity weighted and surface weighted (inset) size distributions obtained from DLS measurements of POPC and POPC/WALP vesicles obtained using different formulation methods: vortexing, extrusion, and sonication.

The average diameters obtained from cumulant analysis for the sonicated vesicles were 0.05–0.30 μm depending on the composition. Please consider that our samples were not centrifugated after sonication. For comparison we measured multilamellar vesicles (vortexed only without sonication nor extrusion) obtaining mean sizes of 3.4 ± 4.0 µm without, and 1.0 ± 0.4 µm with 5 mol% of WALP. After extrusion through filters with 0.1 µm pores, sizes of 0.107 ± 0.003 µm and 0.103 ± 0.002 µm respectively were obtained for the same samples. Mean sizes are summarized in Table S1 and DLS size distributions in Figure S1. Increased peptide content from 0 to 5 mol% resulted in a decrease of the size of the vesicles (from 0.25 to 0.10 µm for POPC, and from 0.30 to 0.10 µm for POPC/Chol), which suggests that the peptide changes the mechanical properties of the proteoliposome membrane.

**Table S1**: Hydrodynamic diameter of liposomes and proteoliposomes obtained from DLS cumulant analysis.

| Sample | Formulation method | WALP peptide content (mol%) | Hydrodynamic diameter, ± SD (µm) |
|---|---|---|---|
| POPC | vortexing[a] | 0 | 3.4 ± 4.0 |
| | | 5 | 1.0 ± 0.4 |
| | extrusion[b] | 0 | 0.107 ± 0.003 |
| | | 5 | 0.103 ± 0.002 |
| | sonication[c] | 0 | 0.235 ± 0.013 |
| | | 1 | 0.27 ± 0.11 |
| | | 2.5 | 0.176 ± 0.011 |
| | | 5 | 0.115 ± 0.010 |
| POPC/Chol | sonication[c] | 0 | 0.298 ± 0.071 |
| | | 5 | 0.096 ± 0.011 |
| DMPC[d] | sonication[c] | 0 | 0.165 ± 0.016 |
| | | 5 | 0.098 ± 0.006 |

Typical names/abbreviations used in the literature: [a] multilamellar vesicles/MLV, [b] large unilamellar vesicles/LUV, [c] small unilamellar vesicles/SUV. [c] DMPC-containing samples were sonicated using different apparatus, thus, cannot be easily compared with the other.

### 2. TIME-DEPENDENT FLUORESCNCE SHIFT

#### 2.1 Method

Time-dependent fluorescence shift (TDFS) experiments were performed as follow.

1. 1 mL of liposome or proteoliposome dispersion was transferred in a 1.5 mL quartz spectroscopic cuvette to a sample holder in TCSPC or steady-state apparatus. Temperature was equilibrated for 15 min and kept constant during the experiment using a circulating water bath.
2. Fluorescence decays were measured using our TCSPC setup for $\lambda_{EX} = 373$ nm (slits width = 8 nm), and $\lambda_{EM} = 400$-540 nm (10 nm step, slits width = 12 nm).
3. Emission spectrum at 390-600 nm (1 nm step) was measure for $\lambda_{EX} = 373$ nm; excitation and emission slits = 1.2 nm.
4. Fluorescence decays were fitted with 3-exponential function using the iterative reconvolution procedure in DAS6 software (IBH, UK).
5. The fitted parameters and the steady-state emission spectrum were used to reconstruct time-resolved emission spectra (TRES), which were then fitted with log-normal function to determine position of their maxima, $\nu(t)$, and their width (full-width and half-maximum, FWHM($t$)) using Matlab (MathWorks) script.
6. $\nu(t)$ was used to determine $\Delta\nu$ and $\tau_R$. These parameters reflect membrane hydration and local lipid mobility, respectively. $\Delta\nu = \nu(0) - \nu(\infty)$, where $\nu(0)$ is TRES position immediately after electronic excitation estimated to be 23 800 $cm^{-1}$ using procedure in ref. (1), and $\nu(\infty)$ is spectrum position after dipolar relaxation of solvent is completed (15 ns was chosen for all samples). Mean integrated relaxation time was calculated as $\tau_R = \int_0^{\infty} (\nu(t) - \nu(\infty))/\Delta\nu \; dt$.
7. A second estimate of the relaxation time was taken as a time at which FWHM(t) was maximum.

#### 2.2 Results

Comparison of TDFS and GP results given in Figure S2, demonstrates parallelism between $GP_{330\text{-}360}$ and TDFS relaxation time, $\tau_R$. which corresponds to the mobility of the hydrated phospholipids in Laurdan vicinity. As discussed in the main text Laurdan is located slightly below the lipid glycerol level and probes predominantly hydrated lipid carbonyls. The total spectral shift, $\Delta\nu$, does not change much for the herein measured samples.

Slightly increased $\Delta\nu$ values were obtained only for POPC/Chol and POPC/Chol/WALP samples measured at 37°C. These differences were still very close to the experimental error, especially when compared with the changes observed for $\tau_R$. Therefore, we conclude that the GP results discussed in this work should be interpreted in terms of lipid fluidity and not hydration.

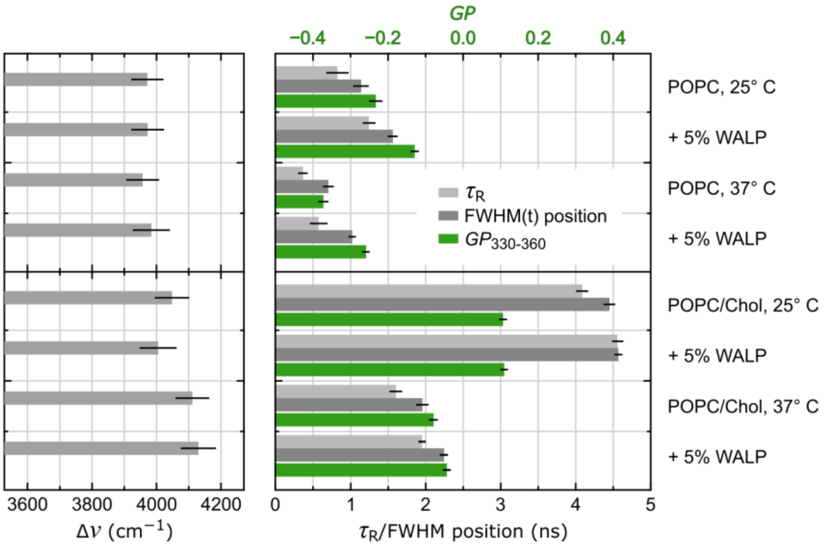


**Figure S2**. Time-Dependent Fluorescence Shift (TDFS) versus Generalized Polarization (GP). (left panel) TDFS total spectral shift, $\Delta\nu$. (right panel) TDFS relaxation time, $\tau_R$, TDFS position of FWHM($t$) maximum (bottom axis), and $GP_{330\text{-}360}$ (top axis). Error bars represent standard deviation.

Figure S2 presents two parameters that reflect the speed of relaxation process. The integrated relaxation time, $\tau_R$, is not always the best measure of the relaxation kinetics. It is especially true for the slow relaxing systems. ”Slow” means here slower than the fluorescent lifetime of the probe. In such case the TRES position does not fully converge to $\nu(\infty)$ value. One possible solution of this issue is to extrapolate $\nu(t)$ values. Another one, is to use the profile of FWHM($t$). It is typical, that during relaxation process TRES change their width. For more precisely defined states, which are the initial Franck-Condon state and the fully dipolarly-relaxed $S_1$ state, TRES are narrower, but in between these extremes they get broadened, which reflects the multitude of intermediate states during the relaxation process. This is reflected in the shape of FWHM($t$) curve with its pronounced maximum during the relaxation process. Position of this maximum corresponds well with the mean relaxation time. In our case both values ($\tau_R$, and the position of FWHM($t$) maximum) are in very good agreement and are only moderately but systematically shifted. $\tau_R$ relaxation times are smaller than those obtained from FWHM($t$) curves, but the relative changes between the samples are preserved.

### 3. FÖRSTER RADIUS CALCULATION

Förster radius was calculated according to:

$$R_0 = 0.0211 \cdot \sqrt[6]{\kappa^2 n^{-4} Q_D \int_0^\infty F_D(\lambda)\varepsilon_A(\lambda)\lambda^4 d\lambda}, \tag{S1}$$

where $\kappa^2$ is an orientation factor assumed to be 2/3, $n$ is the refractive index assumed to be 1.4, $Q_D$ is the quantum yield of the donor in the absence of acceptor taken as 0.12, $F_D$ is donor fluorescence with the total intensity normalized to 1, $\varepsilon_A$ is the extinction coefficient of the acceptor, and λ is a wavelength (27).

### 4. SIMPLIFIED POINT METHOD FOR THE S2 CONTRIBUTION REMOVAL

#### 4.1 Derivation

The method assumes that $L_{S2}$ and $L_{FRET}$ lineshapes are determined properly and do not change within the experiment. We chose two excitation wavelengths for which $L_{S2}$ and $L_{FRET}$ differ significantly, namely, 250 and 280 nm. In the cholesterol-free case, $L_{S2}$(250 nm) = 0.68, $L_{S2}$(280 nm) = 0.44, $L_{FRET}$(250 nm) = 0.21, and $L_{FRET}$(280 nm) = 0.73. Then we substitute them to Eq. 2 separately for 440 and 490 nm obtaining the following set of equations:

$$\begin{cases} F_{250/440} = L_{\mathrm{S2}}(250) \cdot s_{440} + L_{\mathrm{FRET}}(250) \cdot f_{440} \\ F_{280/440} = L_{\mathrm{S2}}(280) \cdot s_{440} + L_{\mathrm{FRET}}(280) \cdot f_{440} \end{cases},$$
$$\begin{cases} F_{250/490} = L_{\mathrm{S2}}(250) \cdot s_{490} + L_{\mathrm{FRET}}(250) \cdot f_{490} \\ F_{280/490} = L_{\mathrm{S2}}(280) \cdot s_{490} + L_{\mathrm{FRET}}(280) \cdot f_{490} \end{cases}. \quad \text{(S2)}$$

$F_{\lambda_{\mathrm{EX}}/\lambda_{\mathrm{EM}}}$ denotes fluorescence intensity measured at an excitation wavelength of $\lambda_{\mathrm{EX}}$ and an emission wavelength of $\lambda_{\mathrm{EM}}$. Eq. S2 can be readily solved to get $s_{440}$, $s_{490}$, $f_{440}$, and $f_{490}$ parameters, which can be then used in Eq. 3 and 4 to finally get:

$$GP_{\mathrm{S2}} = \frac{F_{250/440} - F_{250/490} - \beta\,(F_{280/440} - F_{280/490})}{F_{250/440} + F_{250/490} - \beta\,(F_{280/440} + F_{280/490})},\ \beta = \frac{L_{\mathrm{FRET}}(250\,nm)}{L_{\mathrm{FRET}}(280\,nm)}, \quad \text{(S3)}$$

$$GP_{\mathrm{FRET}} = \frac{F_{250/440} - F_{250/490} - \alpha\,(F_{280/440} - F_{280/490})}{F_{250/440} + F_{250/490} - \alpha\,(F_{280/440} + F_{280/490})},\ \alpha = \frac{L_{\mathrm{S2}}(250\,nm)}{L_{\mathrm{S2}}(280\,nm)}. \quad \text{(S4)}$$

Based on Eq. S3 and S4 the errors of the GP values can be calculated using partial derivative method, namely:

$$\Delta GP = \sum_{i=1}^{n} \left| \frac{\partial GP}{\partial p_i} \right| \Delta p_i, \quad \text{(S5)}$$

where $\Delta GP$ is the estimated error, and $\Delta p_i$ is the error of the $i^{\mathrm{th}}$ parameter ($p_i$). Since the obtained equations are long, we use a more condensed symbols for the variables (see Table S2):

$$\Delta GP_{FRET} = 2[F_{54} + F_{59} - \alpha(F_{84} + F_{89})]^{-2}[|F_{54}F_{89} - F_{59}F_{84}|(\alpha\Delta L_{S8} + \Delta L_{S5}) \ldots + |F_{54} - \alpha F_{84}|(\alpha\Delta F_{89} + \Delta F_{59}) + |F_{59} - \alpha F_{89}|(\alpha\Delta F_{84} + \Delta F_{54})], \quad \text{(S6)}$$

$$\Delta GP_{S2} = 2[F_{54} + F_{59} - \alpha(F_{84} + F_{89})]^{-2}[|F_{54}F_{89} - F_{59}F_{84}|(\alpha\Delta L_{F8} + \Delta L_{F5}) \ldots + |F_{54} - \alpha F_{84}|(\alpha\Delta F_{89} + \Delta F_{59}) + |F_{59} - \alpha F_{89}|(\alpha\Delta F_{84} + \Delta F_{54})]. \quad \text{(S7)}$$

**Table S2**: Symbols for Eq. S6 and S7.

| Symbol | $\lambda_{\mathrm{EX}}$ (nm) | $\lambda_{\mathrm{EM}}$ (nm) | Description |
|---|---|---|---|
| $F_{54}$ | 250 | 440 | Fluorescence intensities measured for the sample of interest |
| $F_{59}$ | 250 | 490 | |
| $F_{84}$ | 280 | 440 | |
| $F_{89}$ | 280 | 490 | |
| $L_{\mathrm{S5}}$ | 250 | 440 & 490 | Values of the excitation spectral line shapes |
| $L_{\mathrm{S8}}$ | 280 | 440 & 490 | |
| $L_{\mathrm{F5}}$ | 250 | 440 & 490 | |
| $L_{\mathrm{F8}}$ | 280 | 440 & 490 | |

#### 4.2 Comparison

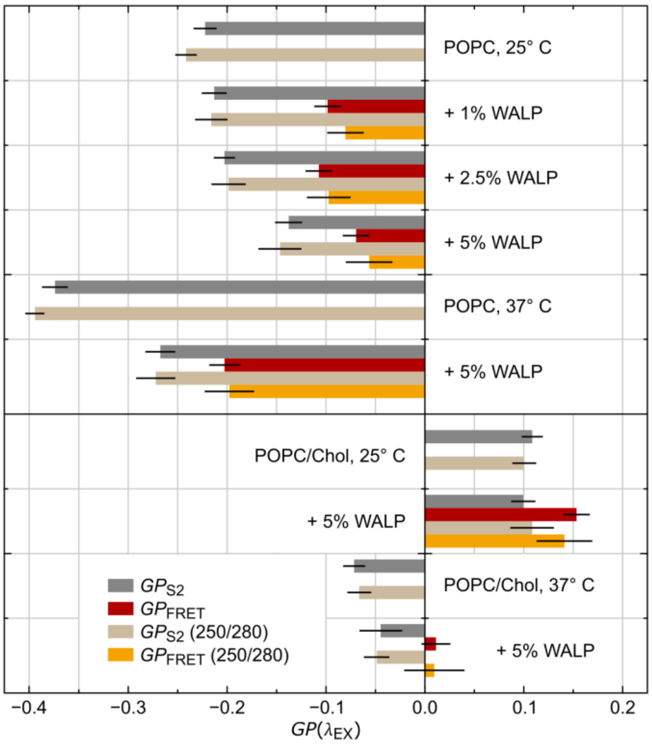


**Figure S3**. Comparison of two methods of FRET-GP analysis: first – based on lineshape fitting, second – based on the values of fluorescence intensities measure for excitation at 250 and 280 nm. GP values obtained with the second method are marked with "(250/280)". All FRET-GP values obtained for different content of WALP incorporated in POPC and POPC/Chol membranes measured at 25°C and 37°C. Error bars represent standard deviation. This figure corresponds to Figure 4H.

### 5. $S_2$ AND FRET LINESHAPES FOR DMPC PHASE TRANSITION

Using the method developed based on our POPC-based samples we prepared excitational lineshapes for DMPC/WALP separately for each temperature. As shown in Figure S4, their shapes vary with the temperature, which is particularly visible for the S2 lineshapes. Variability of the corresponding FRET lineshapes is much smaller. With increasing temperature, when DMPC becomes more fluid, the lineshapes approach the one obtained before for POPC (black dashed line). Apparently, the lineshapes reflect the phase of the lipid and would probably be similar for all fluid-phase lipid bilayers.

For the DMPC case, where we probe across the phase transition, the quality of fitting used in spectral demixing procedure can suffer considerably from using inappropriate lineshapes. We tested this using 1) our POPC-derived lineshapes, 2) average lineshapes derived from all DMPC measurements regardless of the temperatures, and 3) DMPC lineshapes obtained for the matching temperature. The fitting obtained with these lineshapes for the DMPC/WALP spectra measured at 10°C, 20°C, 25°C and 37°C are shown in Figure 4S; panels B, C and D, respectively. Panel B shows that at 10°C the curve fitted using the POPC lineshapes (solid line) considerably deviates from the DMPC data showed as dots. Increasing the temperature improves the fitting quality, which at 25°C is already reasonable. This is understandable, since we are using the lineshapes derived for liquid crystallin $L_\alpha$ phase (POPC at 25°C and 37°C). The deviations are large when DMPC lipid is in its gel $P_\beta$ phase, but at high temperatures when DMPC is fluid the fit quality is similar regardless of the lineshapes used.

The quality of the fits is well visible in the residuals plotted in the lower parts of panels B,C and D. Application of the average DMPC lineshapes (panel C) lowers the residuals slightly compared to those for the POPC lineshapes (panel B), but they still do not match those for the lineshapes for dedicated temperature (panel C). As already mentioned these deviations are important for the lower temperatures, while at 37°C all the lineshapes perform well. When talking about the performance of the spectral demixing we should also consider the $GP_{FRET}$ values obtained as a result. Luckily, even for the cases with worse-quality fitting (the POPC lineshapes applied to DMPC/WALP at 10°C) the obtained results are almost identical with those obtained using the proper – dedicated DMPC lineshapes. We also performed the analysis using the simplified point method described above for the same values used for

POPC ($L_{S2}$(250 nm) = 0.68, $L_{S2}$(280 nm) = 0.44, $L_{FRET}$(250 nm) = 0.21, and $L_{FRET}$(280 nm) = 0.73). Comparison of the $GP_{FRET}$ values obtained using all these methods are given in Table S3.

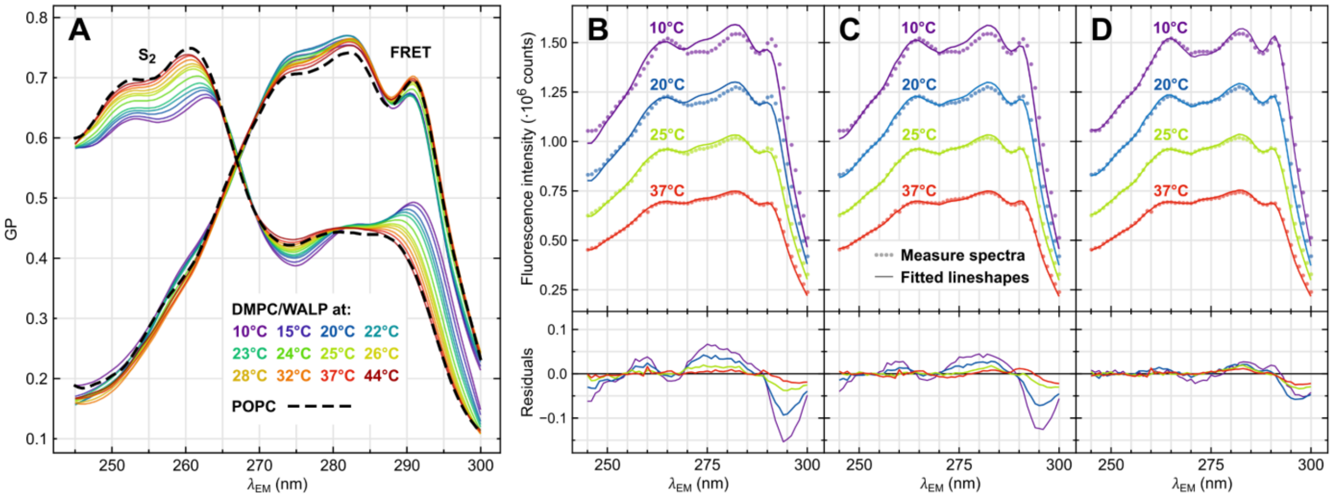


**Figure S4**. Laurdan excitational spectral lineshapes obtained for DMPC/WALP peptidoliposomes at different temperatures and fitting errors they cause in the spectral demixing procedure. (A) Lineshapes for DMPC-based vesicles at different temperatures. The lineshapes obtained for POPC-based samples at 25°C and 37°C is shown as dashed line for comparison. (B-D) Examples of the best fits obtained for the excitation spectra of Laurdan in DMPC/WALP at 10°C, 20°C, 25°C and 37°C for different spectral lineshapes: (B) the POPC-derived lineshapes, (C) averages of all DMPC-derived lineshapes, and (D) DMPC-derived lineshapes for the corresponding temperatures.

**Table S3**: Comparison of $GP_{FRET}$ results obtained for DMPC/WALP peptidoliposomes with different excitational lineshapes and methods.

| Temperature | $GP_{FRET}$[a] | $GP_{FRET}$[b] | $GP_{FRET}$[c] | $\boldsymbol{GP_{FRET}}$[d] |
|---|---|---|---|---|
| 10°C | 0.412 ± 0.012 | 0.408 ± 0.010 | 0.410 ± 0.011 | **0.413 ± 0.011** |
| 20°C | 0.273 ± 0.015 | 0.273 ± 0.015 | 0.274 ± 0.015 | **0.274 ± 0.015** |
| 25°C | 0.145 ± 0.008 | 0.140 ± 0.007 | 0.143 ± 0.007 | **0.142 ± 0.007** |
| 37°C | −0.082 ± 0.009 | −0.087 ± 0.010 | −0.082 ± 0.010 | **−0.086 ± 0.010** |

[a] Excitational lineshapes for POPC / simplified point method for analysis
[b] Excitational lineshapes for POPC / fitting-based spectral demixing method
[c] Average excitational lineshapes for DMPC / fitting-based spectral demixing method
[d] Excitational lineshapes for DMPC matching the temperature / fitting-based spectral demixing method


### 7. APPENDIX S1

Excitation Laurdan spectral lineshapes for $S_2$ and FRET excitations for POPC and POPC/Chol systems

| | POPC | | POPC/Chol | |
|---|---|---|---|---|
| $\lambda$ (nm) | $L_{S2}$ ± SE | $L_{FRET}$ ± SE | $L_{S2}$ ± SE | $L_{FRET}$ ± SE |
| 245 | 0.600 ± 0.009 | 0.188 ± 0.021 | 0.581 ± 0.008 | 0.192 ± 0.007 |
| 246 | 0.605 ± 0.009 | 0.184 ± 0.021 | 0.588 ± 0.008 | 0.196 ± 0.007 |
| 247 | 0.619 ± 0.009 | 0.187 ± 0.021 | 0.599 ± 0.008 | 0.200 ± 0.007 |
| 248 | 0.637 ± 0.009 | 0.192 ± 0.021 | 0.611 ± 0.008 | 0.205 ± 0.007 |
| 249 | 0.659 ± 0.009 | 0.197 ± 0.021 | 0.626 ± 0.007 | 0.213 ± 0.007 |
| 250 | 0.675 ± 0.009 | 0.205 ± 0.020 | 0.642 ± 0.0078 | 0.220 ± 0.007 |

| $\lambda$ (nm) | POPC $L_{S2}$ ± SE | POPC $L_{FRET}$ ± SE | POPC/Chol $L_{S2}$ ± SE | POPC/Chol $L_{FRET}$ ± SE |
|---|---|---|---|---|
| 251 | 0.687 ± 0.009 | 0.212 ± 0.020 | 0.653 ± 0.007 | 0.229 ± 0.007 |
| 252 | 0.695 ± 0.009 | 0.225 ± 0.020 | 0.661 ± 0.007 | 0.244 ± 0.007 |
| 253 | 0.698 ± 0.009 | 0.243 ± 0.019 | 0.666 ± 0.007 | 0.259 ± 0.007 |
| 254 | 0.697 ± 0.009 | 0.263 ± 0.018 | 0.666 ± 0.006 | 0.274 ± 0.007 |
| 255 | 0.698 ± 0.009 | 0.285 ± 0.018 | 0.667 ± 0.006 | 0.294 ± 0.007 |
| 256 | 0.701 ± 0.009 | 0.306 ± 0.017 | 0.669 ± 0.006 | 0.315 ± 0.007 |
| 257 | 0.711 ± 0.009 | 0.325 ± 0.016 | 0.674 ± 0.006 | 0.337 ± 0.007 |
| 258 | 0.725 ± 0.008 | 0.342 ± 0.015 | 0.685 ± 0.006 | 0.355 ± 0.006 |
| 259 | 0.740 ± 0.008 | 0.357 ± 0.014 | 0.696 ± 0.006 | 0.374 ± 0.006 |
| 260 | 0.748 ± 0.008 | 0.374 ± 0.012 | 0.706 ± 0.006 | 0.393 ± 0.006 |
| 261 | 0.749 ± 0.007 | 0.392 ± 0.011 | 0.713 ± 0.006 | 0.411 ± 0.006 |
| 262 | 0.739 ± 0.007 | 0.415 ± 0.010 | 0.714 ± 0.005 | 0.431 ± 0.005 |
| 263 | 0.717 ± 0.007 | 0.445 ± 0.010 | 0.706 ± 0.005 | 0.452 ± 0.005 |
| 264 | 0.685 ± 0.006 | 0.475 ± 0.009 | 0.688 ± 0.005 | 0.476 ± 0.005 |
| 265 | 0.646 ± 0.006 | 0.509 ± 0.009 | 0.661 ± 0.005 | 0.504 ± 0.004 |
| 266 | 0.602 ± 0.006 | 0.539 ± 0.008 | 0.625 ± 0.005 | 0.533 ± 0.004 |
| 267 | 0.559 ± 0.006 | 0.567 ± 0.008 | 0.584 ± 0.005 | 0.564 ± 0.004 |
| 268 | 0.520 ± 0.005 | 0.591 ± 0.009 | 0.542 ± 0.005 | 0.591 ± 0.004 |
| 269 | 0.488 ± 0.005 | 0.617 ± 0.009 | 0.503 ± 0.005 | 0.617 ± 0.004 |
| 270 | 0.463 ± 0.006 | 0.640 ± 0.010 | 0.472 ± 0.005 | 0.644 ± 0.004 |
| 271 | 0.445 ± 0.006 | 0.664 ± 0.010 | 0.450 ± 0.004 | 0.669 ± 0.004 |
| 272 | 0.433 ± 0.006 | 0.684 ± 0.011 | 0.435 ± 0.004 | 0.690 ± 0.004 |
| 273 | 0.425 ± 0.006 | 0.698 ± 0.012 | 0.426 ± 0.004 | 0.705 ± 0.004 |
| 274 | 0.422 ± 0.006 | 0.705 ± 0.013 | 0.419 ± 0.004 | 0.713 ± 0.004 |
| 275 | 0.423 ± 0.007 | 0.707 ± 0.014 | 0.418 ± 0.004 | 0.714 ± 0.004 |
| 276 | 0.427 ± 0.007 | 0.708 ± 0.015 | 0.421 ± 0.004 | 0.716 ± 0.004 |
| 277 | 0.433 ± 0.007 | 0.711 ± 0.016 | 0.428 ± 0.004 | 0.719 ± 0.004 |
| 278 | 0.437 ± 0.007 | 0.716 ± 0.016 | 0.435 ± 0.004 | 0.724 ± 0.005 |
| 279 | 0.441 ± 0.007 | 0.723 ± 0.017 | 0.442 ± 0.004 | 0.730 ± 0.005 |
| 280 | 0.443 ± 0.007 | 0.732 ± 0.017 | 0.447 ± 0.005 | 0.738 ± 0.005 |
| 281 | 0.444 ± 0.007 | 0.739 ± 0.017 | 0.449 ± 0.005 | 0.742 ± 0.005 |
| 282 | 0.443 ± 0.007 | 0.742 ± 0.017 | 0.449 ± 0.005 | 0.747 ± 0.005 |
| 283 | 0.442 ± 0.007 | 0.740 ± 0.016 | 0.449 ± 0.005 | 0.745 ± 0.005 |
| 284 | 0.440 ± 0.007 | 0.726 ± 0.016 | 0.449 ± 0.005 | 0.732 ± 0.006 |
| 285 | 0.440 ± 0.007 | 0.704 ± 0.015 | 0.450 ± 0.006 | 0.707 ± 0.006 |
| 286 | 0.440 ± 0.007 | 0.678 ± 0.014 | 0.453 ± 0.006 | 0.676 ± 0.006 |
| 287 | 0.437 ± 0.007 | 0.658 ± 0.013 | 0.456 ± 0.006 | 0.648 ± 0.006 |
| 288 | 0.431 ± 0.008 | 0.654 ± 0.012 | 0.458 ± 0.006 | 0.636 ± 0.006 |
| 289 | 0.421 ± 0.008 | 0.669 ± 0.011 | 0.458 ± 0.007 | 0.647 ± 0.006 |
| 290 | 0.402 ± 0.009 | 0.688 ± 0.010 | 0.450 ± 0.007 | 0.665 ± 0.006 |
| 291 | 0.378 ± 0.009 | 0.696 ± 0.009 | 0.431 ± 0.007 | 0.672 ± 0.006 |
| 292 | 0.346 ± 0.010 | 0.675 ± 0.008 | 0.403 ± 0.008 | 0.651 ± 0.006 |
| 293 | 0.309 ± 0.011 | 0.624 ± 0.007 | 0.363 ± 0.008 | 0.599 ± 0.006 |
| 294 | 0.268 ± 0.011 | 0.557 ± 0.007 | 0.315 ± 0.008 | 0.528 ± 0.006 |
| 295 | 0.230 ± 0.012 | 0.488 ± 0.006 | 0.267 ± 0.008 | 0.458 ± 0.006 |
| 296 | 0.195 ± 0.012 | 0.427 ± 0.006 | 0.222 ± 0.008 | 0.396 ± 0.005 |
| 297 | 0.166 ± 0.012 | 0.375 ± 0.006 | 0.183 ± 0.008 | 0.344 ± 0.005 |
| 298 | 0.143 ± 0.013 | 0.326 ± 0.006 | 0.151 ± 0.008 | 0.300 ± 0.005 |
| 299 | 0.125 ± 0.013 | 0.280 ± 0.005 | 0.125 ± 0.008 | 0.257 ± 0.005 |
| 300 | 0.110 ± 0.013 | 0.233 ± 0.006 | 0.102 ± 0.008 | 0.215 ± 0.006 |
